# Alpha oscillation, criticality, and responsiveness in complex brain networks

**DOI:** 10.1101/529982

**Authors:** MinKyung Kim, UnCheol Lee

## Abstract

Brains in sleep, anesthesia, and traumatic injury are characterized by significantly limited responsiveness to stimuli. Even during conscious wakefulness, responsiveness is highly dependent on on-going brain activity, specifically, of alpha oscillations (∼10Hz). However, despite many empirical studies, the ways in which specific alpha oscillations induce a large or small response to stimuli have not been elucidated. We hypothesized that the variety of responses to sensory stimuli result from the interaction between state-specific and transient alpha oscillations and stimuli. To justify this hypothesis, we simulated various alpha oscillations in the human brain network, modulating network criticality (a balanced state between order and disorder), and investigated specific alpha oscillation properties (instantaneous amplitude, phase, and global synchronization) that induce a large or small response. We found that near a critical state, a large, complex response is induced when a stimulus is given to globally desynchronized and low-amplitude alpha oscillations, and we also found specific phases of alpha oscillation that barely respond to stimuli. These results imply the presence of temporal windows in the alpha cycle for a large or small response to external stimuli, which is consistent with the periodic perceptual binding at alpha frequency found in empirical studies.

## Introduction

The brain’s responsiveness to external stimuli is significantly dependent on its state. Previous studies with transcranial magnetic stimulation (TMS) have demonstrated that stimulation effects during sleep (NREM), anesthesia, and vegetative states are simple and constrained to local areas, whereas they are more complex and diffuse in the conscious state [1–4]. In the conscious state, perceptual responses to sensory stimuli are associated with ongoing brain activity at the moment when the stimulus is applied. In visual perception tasks, for example, target detectability is dependent on the power and phase of ongoing alpha oscillations (∼10Hz) [5–14]. Recent repetitive TMS (rTMS) studies have suggested that the instantaneous phase of the sensorimotor *μ*-rhythm in the alpha band (8-12Hz) is an important factor in determining the stimulation effect [15, 16].

However, despite many relevant experimental studies, researchers have not elucidated how sensory stimulation interacts with specific amplitudes and phases of alpha oscillation in ways that result in a large or small perceptual response. In this study, we hypothesized that perceptual binding associates with the brain’s network response, and the network response largely depends on the ongoing coupled oscillation properties at stimulus onset. Recent studies have revealed that alpha oscillations transfer information through traveling waves in the cortex, and the global functional connectivity of alpha oscillations reflect conscious and unconscious states [17–24]. Therefore, we assumed that the propagation and complexity of perturbed responses in the brain network may be determined by the alpha oscillations at stimulus onset. In this modeling study, we examined the specific amplitude, phase, and synchronization level of alpha oscillations that evoke a large or small network response.

To identify such a condition, we constructed a large-scale human brain network model with the Stuart-Landau oscillators of the alpha band frequencies (∼10Hz) that are coupled with each other in an anatomically informed human brain connection structure. We tested three dynamic brain states, modulating criticality (below, near, and above critical state). The three brain states produced various transient alpha oscillations in terms of global synchronization, amplitude and phase. The critical state was defined with the largest pair correlation function (PCF), measuring the variance of global synchronization fluctuation across time [25], which is equivalent to susceptibility in statistical physics models. Theoretically, near the critical point, a large dynamic fluctuation would be expected. [26, 27]. Empirically, the brain during conscious wakefulness presents a large dynamic fluctuation, especially in synchronization [28, 29]; however, various brain perturbations (e.g., sleep, anesthesia, traumatic injuries) mitigate this fluctuation, which implies that the brain deviates from criticality [30].

The various alpha oscillations of the three brain states were decomposed into basic oscillation properties: instantaneous global synchronization, amplitude, and phase. We applied a pulsatile stimulus to the brain network in order to investigate the relationship between these alpha oscillation properties at stimulus onset and brain network responsiveness. We found specific phases of alpha oscillation that barely respond to the pulsatile stimulus, which suggests the presence of periodic temporal windows in the alpha frequency cycle for a large/small responsiveness, and which is consistent with the empirical studies [5,6,31–34]. A schematic diagram of the study design is presented in Fig 1.

**Fig 1.**
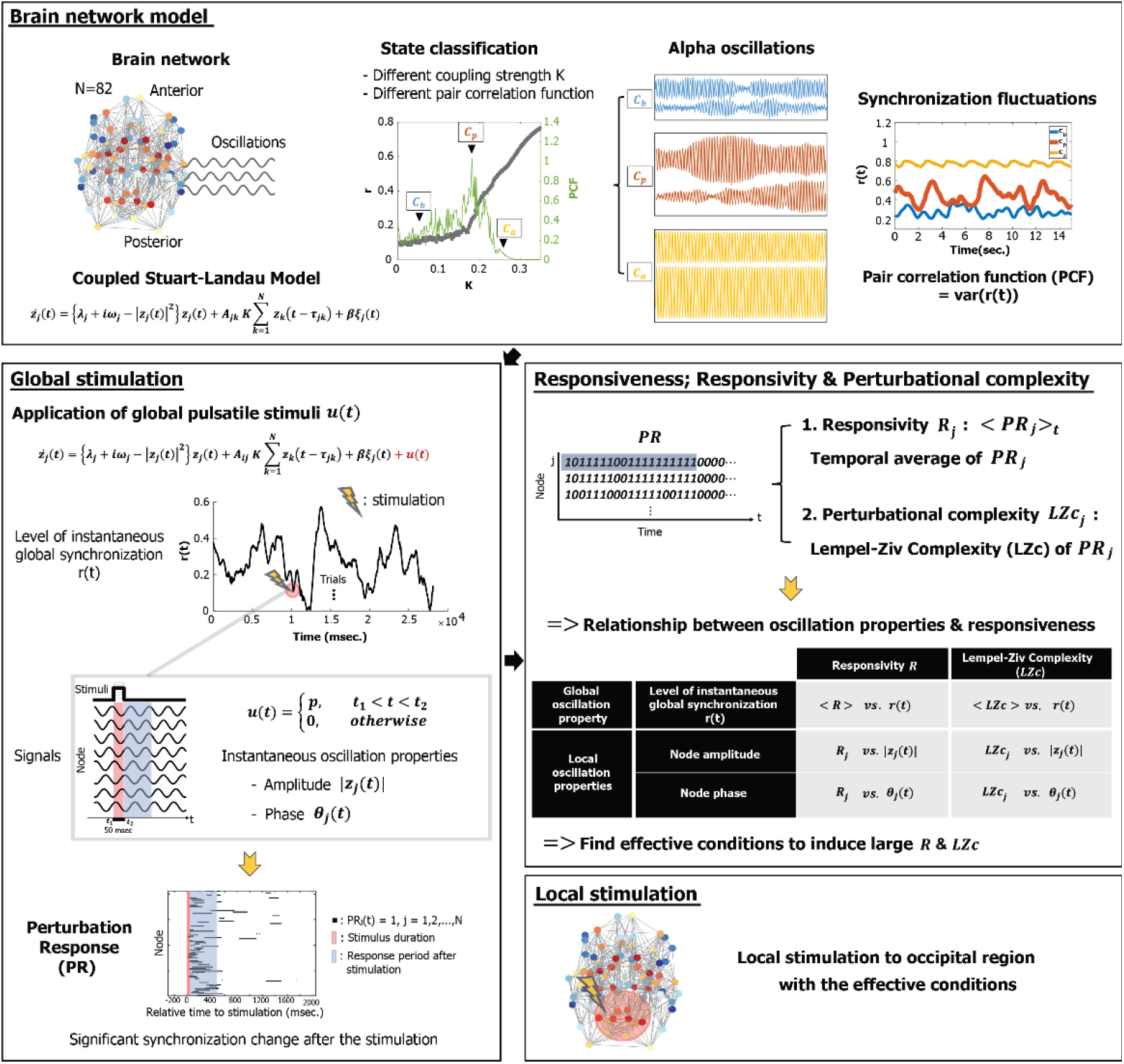
Schematic diagram of the study design. In this study, we simulated alpha oscillations, applying a coupled Stuart-Landau model to a human brain network structure consisting of 82 brain regions. Brain states were classified as below, near, or above a critical state based on the pair correlation function (PCF, the variance of the instantaneous global synchronization level). The critical state *C_p_* was defined with the largest PCF. Global pulsatile stimuli *u* were applied to the 82 alpha oscillation nodes in three brain states, and the following perturbation responses of phase synchronization (*PR_j_*(*t*)) were binarized by comparing them to the pre-stimulus period. Using *PR_j_*, we first defined a responsivity *R_j_*, an average of *PR_j_* during certain epochs after stimulation, to measure the magnitude of the response. Second, the Lempel-Ziv complexity (LZc) of *PR_j_* was calculated to measure the perturbational complexity of the response. Then we investigated the relationship between the ongoing alpha oscillation properties (global synchronization, amplitude, and phase) at stimulus onset and brain network responsiveness to find specific alpha oscillation conditions that induce large (or small) responsiveness. Finally, we stimulated the occipital region and compared effective, random, and less effective stimulation conditions.

## Results

### Alpha oscillations near and far from a critical state

Various alpha oscillations in the human brain network were simulated in three brain states, *C_b_*, *C_p_*, and *C_a_* (below, near, and above a critical state, see “Materials and Methods” for details). To simplify our model, we applied a simple form of stimulus (a pulsatile stimulus) to all nodes for 50 msec. In total, 600 stimulation trials were performed for each state. First, we examined the characteristics of alpha oscillations near and far from a critical state. Fig 2A provides examples of distinct global synchronization fluctuations for the three states. Synchronization fluctuation was measured with the pair correlation function (PCF). Figure 2B presents the average synchronization <*r*(*t*)> and PCF of the alpha oscillations at *C_b_*, *C_p_*, and *C_a_*. Twenty different frequency configurations of the brain network were tested. The average levels of synchronization at *C_p_* (orange triangle) are widely distributed between the small and large average levels of synchronization at *C_b_* (blue circle) and *C_a_* (yellow square). The alpha oscillations at *C_p_* (orange triangle) show a larger PCF (Mean±SD = 1.39±0.68) than those of *C_b_* (blue circle) and *C_a_* (yellow square), which suggests a large variance of brain network dynamics at C_p_. Figure 2C and D demonstrate the instantaneous synchronization and instantaneous amplitude of the alpha oscillations at stimulus onset. At *C_p_* (orange), both the instantaneous synchronization levels *r_s_* and amplitudes |*Z_s_*| at the 600 stimulus onsets are variable between the two small and large *r_s_* and |*Z_s_*| of *C_b_* (blue) and *C_a_* (yellow), respectively (In Fig 2C, Median±SD, 0.11±0.04 for *C_b_*, 0.32±0.14 for *C_p_*, and 0.78±0.05 for *C_a_*; in Fig 2D, 0.21±0.04 for *C_b_*, 0.48±0.57 for *C_p_*, and 1.75±1.65 for *C_a_*). In the next sections, we will show how these state-specific alpha oscillations respond to the same stimulus and identify specific conditions of alpha oscillations that produce a large or small response in the brain network.

**Fig 2.**
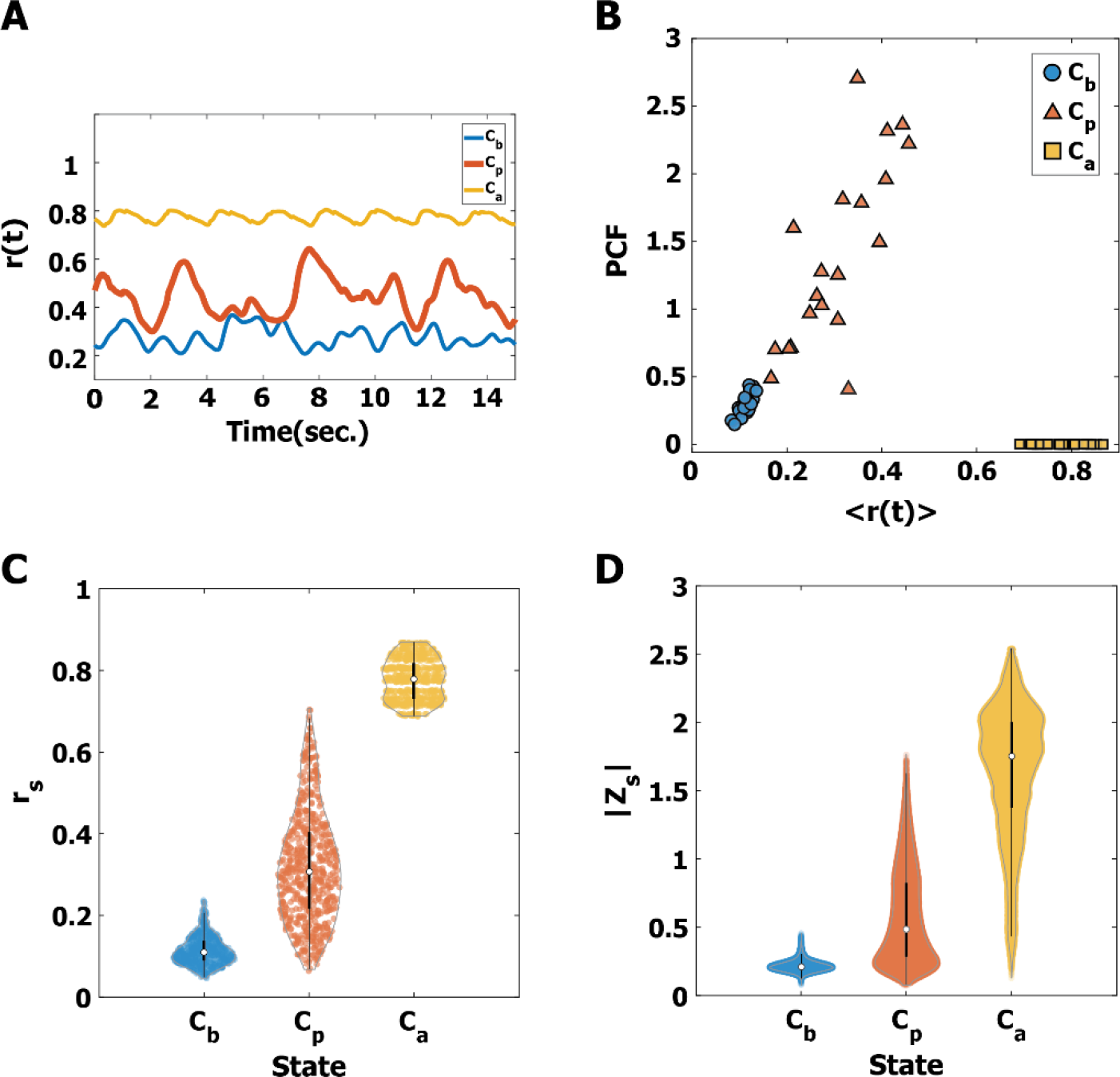
Characteristics of alpha oscillations near and far from a critical state, *C_b_*, *C_p_*, **and** *C_a_*. (A) Examples of the global synchronization fluctuations at *C_b_* (blue), *C_p_* (red), and *C_a_* (yellow) for one initial frequency configuration in the brain network. (B) The average instantaneous global synchronization *r*(*t*) and the pair correlation functions (PCF) for *C_b_* (blue), *C_p_* (red), and *C_a_* (yellow) for 20 different initial frequency configurations in the brain network. Without a stimulus, the critical state *C_p_* is characterized by the largest PCF (Mean±SD = 1.39±0.68), compared to the PCF of *C_b_* and *C_a_*. (C) The estimated density distributions of instantaneous global synchronization (*r_s_*) at the 600 stimulus onsets for *C_b_* (blue), *C_p_* (red), and *C_a_* (yellow). The small white circles indicate the median of *r_s_* and the thick black lines indicate standard deviation. The estimated Gaussian density distribution for the 600 stimuli is plotted with a gray line. The alpha oscillations at *C_p_* have a broad and balanced distribution (Median±SD = 0.32±0.14). (C) The estimated density distributions of the alpha oscillation amplitudes |*Z_s_*| at the 600 stimulus onsets for *C_b_*, *C_p_*, and *C_a_*. The |*Z_s_*| values at *C_b_* and *C_a_* are biased with small and large amplitudes, respectively, while the |*Z_s_*| at *C_p_* is relatively balanced between the others. The distinct alpha oscillation properties at stimulus onset may result in different responses of the brain network to the stimulus.

### Distinct responses of the alpha oscillations near and far from a critical state

Alpha oscillations at diverse states of criticality show distinct responses to the stimulus. Fig 3A shows examples of the instantaneous global synchronization level *r*(*t*) at *C_b_*, *C_p_*, and *C_a_*. Each triangle indicates a discrete application of the stimulus. One pulsatile stimulus was applied at random per simulation; a total of 600 such simulations were performed for each state. To calculate the responsiveness of each stimulation trial, we first defined a perturbation response (PR) of phase synchronization (*PR_j_*(*t*)) for node *j* (j = 1, 2, …, 82) at time t, setting *PR_j_*(*t*) as 1, if the synchronization value of node *j* was significantly different from the pre-stimulus period (one second); otherwise *PR_j_*(*t*) was set as 0 (see “Materials and Methods” for details). Fig 3B presents examples of PR. The black dot represents *PR_j_*(*t*) equals 1 and the red and blue shaded areas indicate the pulsatile stimulus of 50 msec and the response period, respectively. The average *PR_j_*(*t*) was calculated over 600 stimulation trials. Fig 3C shows the temporal changes of the median (thick line) and of the 25% and 75% quantiles (thin lines) of the average PR for the three states. The results show that the PR at *C_p_* persists longer than the PRs at *C_b_* and *C_a_* (sign test, p<0.05 after 190 msec, Fig 2C). We used the PRs during the first 500 msec to quantify responsiveness for further analysis. In addition to the PR of phase synchronization, we also calculated the PR of the amplitudes of the alpha oscillations. These results are presented in the supplementary materials.

**Fig 3.**
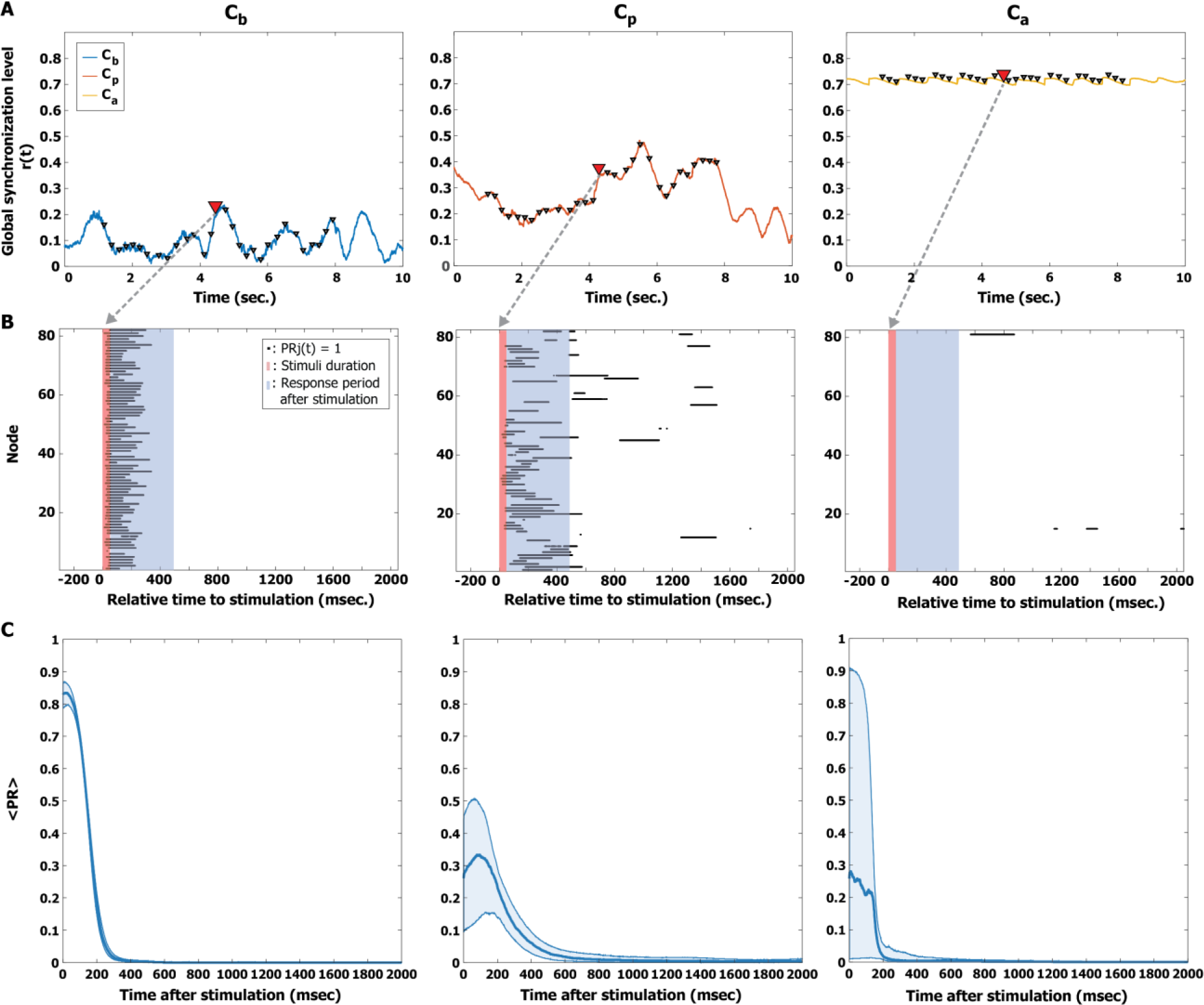
Quantifying the perturbation responses (PRs). (A) Examples of the global synchronization level *r*(*t*) for the three brain states, *C_b_* (left), *C_p_* (middle), and *C_a_* (right). A triangle indicates the timing of a stimulus. The response to a single pulsatile stimulus was stimulated repeatedly 600 times with random stimulus timings. (B) The PR of a pulsatile stimulus is presented for the three brain states. The red and blue shaded areas indicate the stimulus period (50 msec) and response period (500 msec), respectively. The black dot at each node indicates the time point at which phase synchronization significantly increased after the stimulus; *PR_j_* = 1. (C) The average *PR_j_*(*t*) was calculated over 600 stimulation trials, using a new time axis in which the timing of the stimulus coincides with zero. The thick lines indicate the median of the average *PR* and the shaded areas cover the 25% to 75% quantiles. The *PR* persists longer at *C_p_* than at *C_b_* and *C_a_* (sign test, p<0.05 after 190 msec).

### Magnitude and complexity of perturbation responses

In this study, we defined two responsiveness indexes to quantify the PRs. First, we defined the responsivity *R_j_* of a node *j*, counting the total amount of 1s in *PR_j_* (significantly changed phase synchronization) during the first 500 msec after the stimulus (see “Materials and Methods” for details), which measures the magnitude of perturbation responses. We also defined the Lempel-Ziv Complexity (LZc) [35] of the *PR_j_* for a node j, which measures the perturbational complexity (see “Materials and Methods” for details). The LZc measures the amount of information contained in the spatiotemporal pattern of PR, which is similar to the Perturbational Complexity Index (PCI) that has been successfully applied to quantify the level of consciousness in sleep, anesthesia, and pathologic unconsciousness [2, 4].

### Correlation between responsiveness and synchronization of alpha oscillations

First, we investigated the relationship between instantaneous synchronization at stimulus onset (*r_s_*, *s* = 1,2, …, 600) and the responsiveness indexes. The average responsivity < *R* > was calculated by taking the average of *R_j_* over all nodes. Fig 4A presents the Spearman correlation coefficients of the three states (*C_b_*, *C_p_*, and *C_a_*) between <R> and the instantaneous synchronization *r_s_* for 600 stimulation trials. Only at *C_p_*, < *R* > shows a significant negative correlation with *r_s_* (Spearman correlation *ρ_R_* = −0.48, *p* < 0.001 for *C_p_*), suggesting a more desynchronized alpha oscillation induces a larger magnitude of perturbation response at *C_p_*.

**Fig 4.**
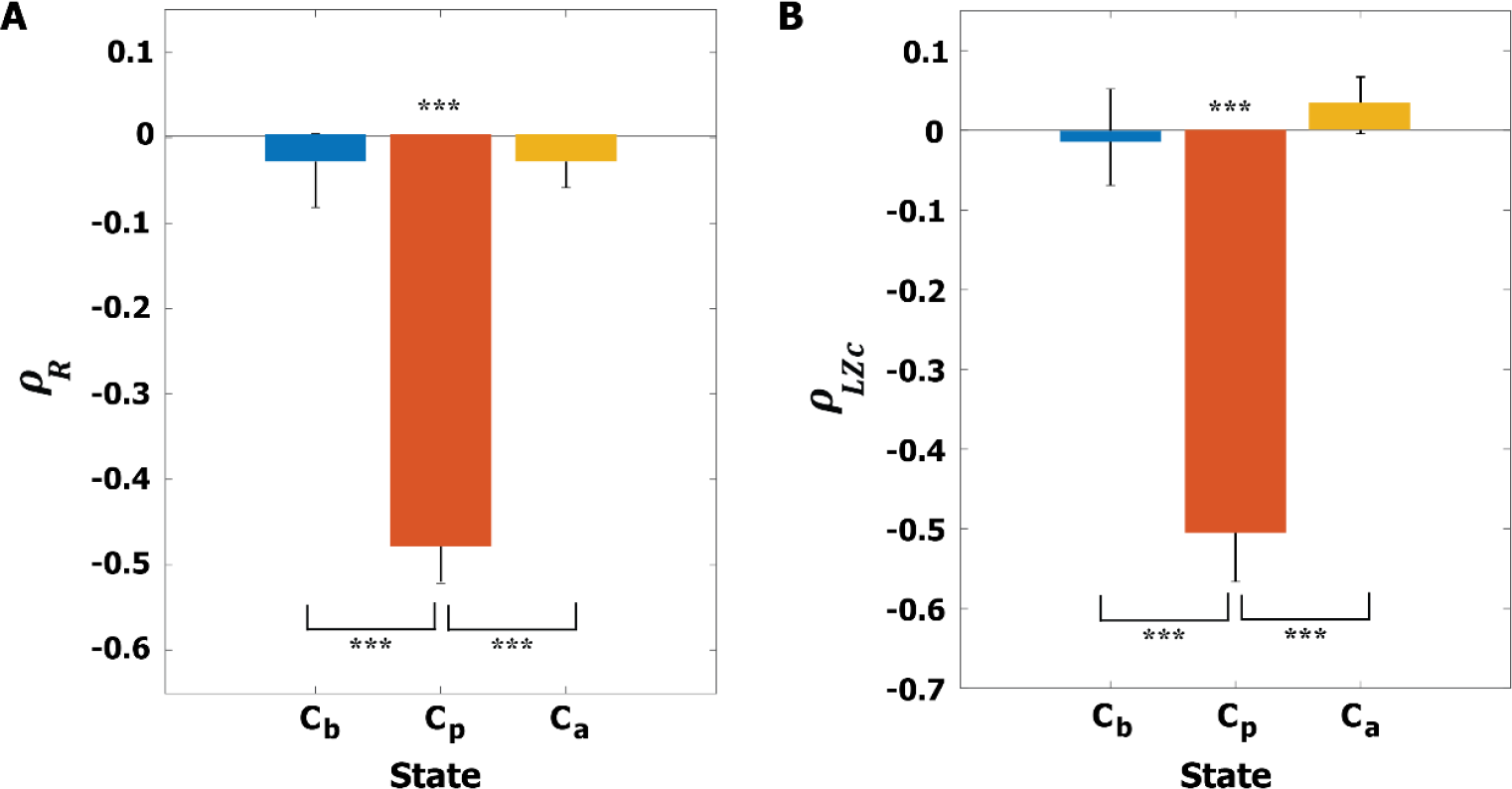
Correlations between instantaneous synchronization (*r_s_*) at stimulus onset and the responsiveness of the brain network. (A) Spearman correlation coefficient *ρ_R_* between *r_s_* and the average responsivity <R> and (B) Spearman correlation coefficient *ρ_LZc_* between *r_s_* and the average complexity of perturbation response <LZc> for *C_b_* (blue), *C_p_* (red), and *C_a_* (yellow). The Spearman correlations were calculated across 600 stimulation trials. The average Spearman correlation coefficients at *C_p_* for both responsiveness indexes are −0.49 and −0.51, respectively, with the significance level ***p<0.001. At *C_p_*, applying a stimulus at a lower level of global synchronization induces a larger magnitude and complexity of perturbation response. A Kuruskal-Wallis test with a Tuckey-Kramer multiple comparison test was performed to compare the correlation coefficients across the three states (***p<0.001).

Next, we calculated the LZc at each time point and averaged it over 500 msec, defining < *LZc* > (See “Materials and Methods” for details). Fig 4B presents the Spearman correlation coefficients between *r_s_* and < *LZc* > for the three brain states. Only at *C_p_*, < *LZc* > is significantly correlated with *r_s_* (Spearman correlation *ρ_LZc_* = −0.52, *p* < 0.001 for *C_p_*). At *C_p_*, a more desynchronized alpha oscillation induces a larger complexity of perturbation response and the complexity itself is the largest among the three states (Fig S1, mean±SD = 0.57±0.05, 0.67±0.17, and 0.39±0.18 for *C_b_*, *C_p_*, and *C_a_*).

We found that near a critical state, stimulating a globally more desynchronized alpha oscillation (small *r_s_*) gave rise to greater brain responsiveness in terms of both magnitude and complexity of perturbation response, whereas far from a critical state, responsiveness was relatively small and not correlated with the global synchronization of alpha oscillations.

### Specific phases of alpha oscillation unresponsive to stimulus

In the previous section, we found a significant correlation between the synchronization of alpha oscillations and responsiveness. In this section, we focus on the critical state *C_p_* and explore whether the amplitude and phase of alpha oscillation also play a role in responsiveness. Interestingly, we found specific phases of alpha oscillation that barely respond to stimuli. We first classified all nodes into two groups according to their oscillation properties—*r_s_* (instantaneous global synchronization level) and |*Z_s_*| (instantaneous amplitude)—at stimulus onset. These nodes were classified into high/low *r_s_* (HS/LS) and high/low |*Z_s_*| (HA/LA) by comparing them to the average *r_s_* and |*Z_s_*|, respectively. We further separated the phases of alpha oscillation into 30 phase bins of *12°* intervals. Fig 5A and B present responsiveness (R and LZc) in each phase bin for the low synchronization and low amplitude (LS & LA) group and high synchronization and high amplitude (HS & HA) group. The thick lines indicate the median value of responsiveness for each phase bin and the shaded area covers the 25% to 75% quantiles of the values of each phase bin for the 600 stimulation trials. We found that average responsiveness of LS & LA is larger than that of HS & HA (p<0.001, A Kruskal-Wallis test). Notably, the HS & HA group showed significant phase dependence. For the purpose of statistical evaluation, we separated these phases into two phase regimes, 60° to 240° and 240° to 60°. The HS & HA group showed significant phase dependence in both responsiveness measures (R and LZc) (Wilcoxon rank sum test, Fig 5C); in particular, stimuli applied to specific phases from 60° to 240° failed to evoke significant brain response (***p<0.001). These results suggest the existence of a periodic temporal window that does not significantly respond to external stimuli. We also tested various other conditions: LS & HA, HS & LA, different stimulus strengths, and the two other states, *C_b_* and *C_a_* (See the S2 Fig for LS & HA and HS & LA; the S3– S7 Figs for the responses to different stimulus strengths; and S8 Fig and S9 Fig for the two other states, *C_b_* and *C_a_*). An excessively small stimulus failed to evoke a response, whereas an excessively strong stimulus produced large responses that eliminated the difference between LS & LA and HS & HA. Phase dependence was also observed in *C_b_* and *C_a_*; however, in this study we focused on the response behavior at critical state *C_p_* to interpret the empirical results during conscious wakefulness.

**Fig 5.**
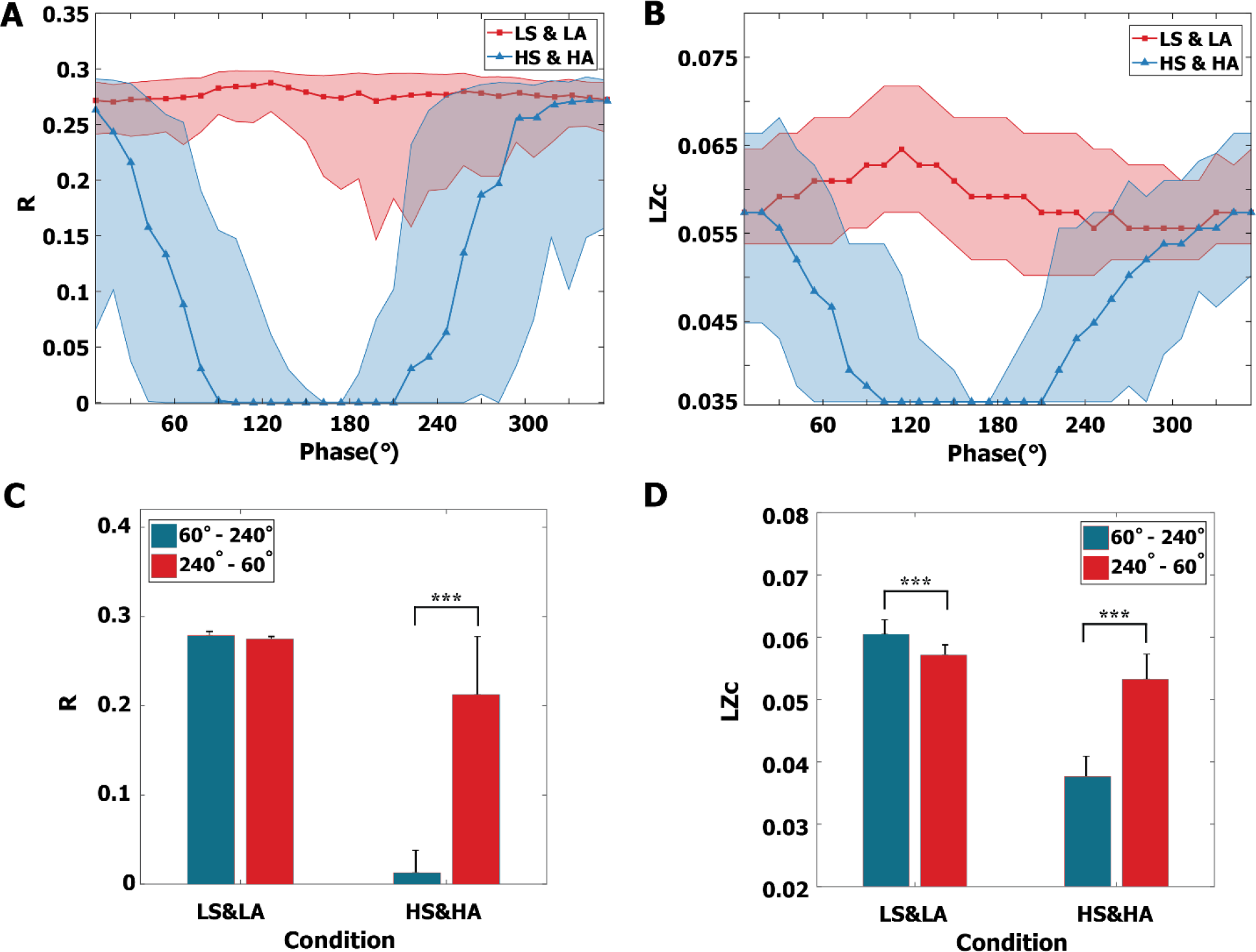
Phase dependence of responsiveness at *C_p_*. The instantaneous alpha oscillations at stimulus onset were separated into two groups: low synchronization and low amplitude (LS & LA) and high synchronization and high amplitude (HS & HA). The phases of alpha oscillation were segmented into 30 phase bins of 12° intervals. (A) The responsivity *R* and (B) the complexity *LZc* of perturbation responses for the HS & HA group significantly depend on the phase of alpha oscillation. The *R* and *LZc* of specific phases (around 180°) show a lower responsiveness to stimuli. A thick red (blue) line indicates the median value; the shaded areas cover the 25% to 75% quantiles. (C and D) For statistical evaluation, the phases of alpha oscillation were separated into two regimes (60°-240° and 240°-60°). In the LS & LA group, stimulation induced greater responsiveness than that in the HS & HA group for both (C) *R* and (D) *LZc* (p<0.001, multiple comparison test using the Tukey-Kramer method). Phase dependence is more prominent in the HS & HA condition than the LS & LA condition, showing larger responsiveness (R and LZc) in the 60°-240° regime (***p<0.001, Wilcoxon rank sum test). In the case of high-amplitude, highly synchronized alpha oscillations, the results imply the existence of temporal windows (with an alpha frequency cycle) that barely respond to external stimuli. Error bar indicates standard deviation.

### Testing global stimulus conditions with a local stimulus

In the previous sections, we examined the various degrees of responsiveness of alpha oscillations to a global stimulus. For this section, we tested if stimulation conditions for large/small responsiveness to the global stimulus also hold for a local stimulus. Here we applied a pulsatile stimulus to nodes within a radius of 50 mm centered on the left cuneus in the occipital region to roughly simulate stimulation of the visual area (13 regions; (left) cuneus, inferior parietal, isthmus cingulate, lateral occipital, lingual, pericalcarine, precuneus, superior parietal, (right) cuneus, isthmus cingulate, lingual, pericalcarine, precuneus). We considered three stimulation conditions: effective, non-effective, and random stimulation. The effective stimulation condition was defined as the phases from 240° to 60° of LS & LA (red bar in LS & LA of Fig 5C & 5D) that induce a large degree of responsiveness. The non-effective stimulation conditions were defined as the phases from 60° to 240° of HS & HA (green bar in HS & HA of Fig 5C & 5D) that induce a small degree of responsiveness. For the random stimulation condition, we randomly selected the stimulus onset, which may correspond to conventional stimulation methods. In comparisons of responsiveness (*R* and *LZc*), we found that the effective (less effective) stimulation condition induces larger (smaller) responsiveness (Fig 6A and 6B), while the responsiveness of the random stimulation condition fell in the middle. The R and LZc of the perturbation responses were significantly different across the three conditions (Kruskal-Wallis test, p<0.001; in addition, a multi-comparison test was performed using the Tukey-Kramer method for R and LZc, respectively. **p<0.005 and ***p<0.001). The results demonstrated that the effective and less effective stimulation conditions we identified for global stimulation can be also applied to local stimulation.

**Fig 6.**
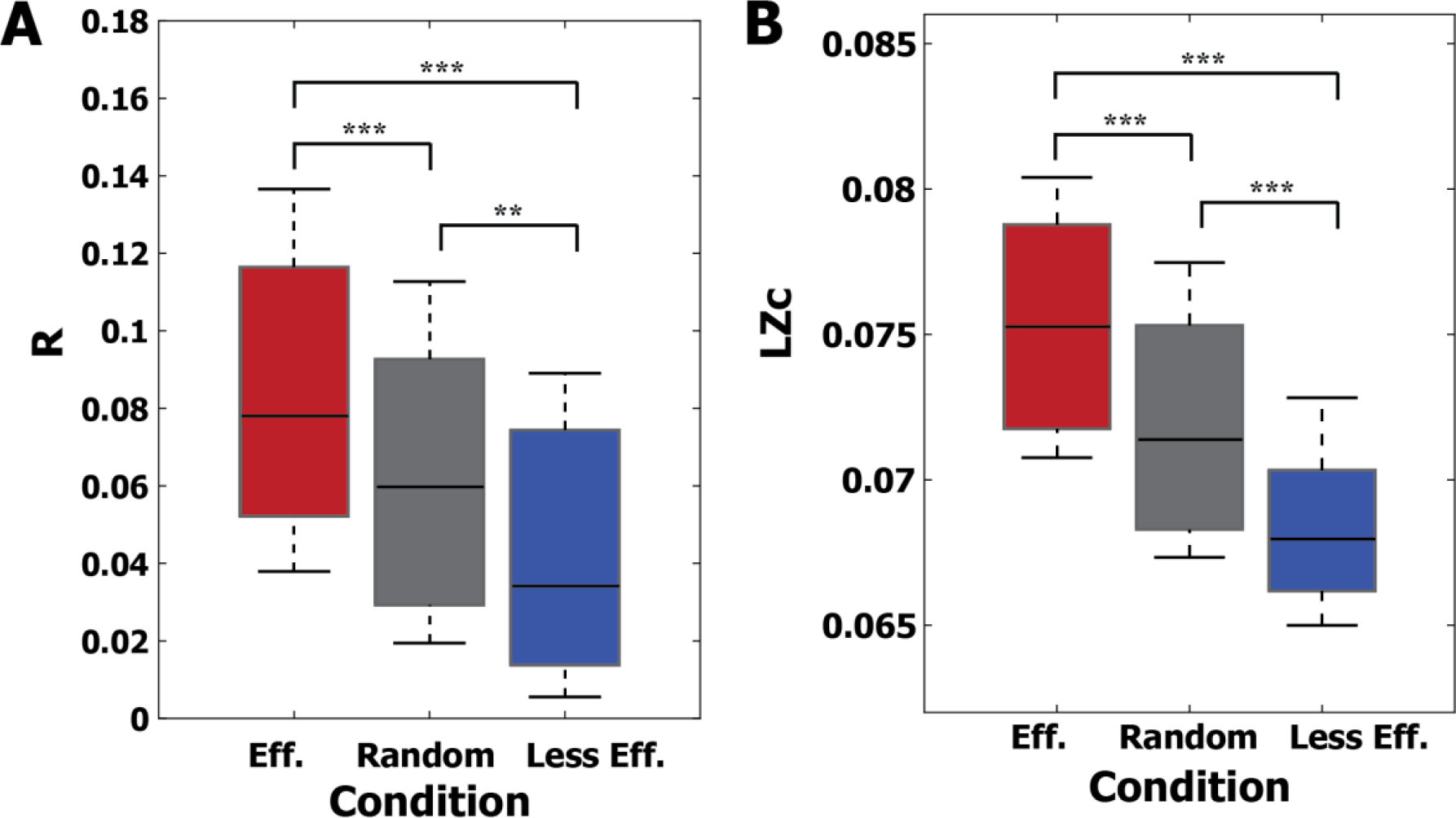
Testing the responsiveness of a local stimulus under (effective, random, and less effective) global stimulation conditions. Three global stimulation conditions were tested to see if they also hold for local stimulation in the brain network. (A) *R* and (B) *LZc* of the occipital region stimulation were compared under three global stimulation conditions. Under the effective stimulation condition, specific phases of alpha oscillation (240° to 60°) were stimulated with low synchronization and low amplitude (red). Under the random stimulation condition, stimulation of alpha oscillations was random (gray). In the less effective stimulation condition, specific phases of alpha oscillation (60°-240°) were stimulated with high synchronization and high amplitude (blue). For occipital region stimulation, the effective (less effective) stimulation conditions produce larger (smaller) responsiveness. The responsiveness of the three conditions are significantly different in both *R* and *LZc* (Kruskal-Wallis test, p<0.001). Multiple comparison tests were performed with the Tukey-Kramer method (**p<0.005 and ***p<0.001). The results showed that the effective and less effective stimulation conditions of global stimulation also hold for local stimulation.

## Discussion

Empirical studies have suggested that brain responsiveness is associated with ongoing brain activities at specific frequencies (e.g., alpha oscillation, ∼10Hz) when a stimulus is applied. In this study, we hypothesized that brain responsiveness is determined by the interaction between a stimulus and coupled alpha oscillations in the brain network. We simulated various alpha oscillations for three brain states (*C_b_*, *C_p_*, and *C_a_*) and investigated the conditions of alpha oscillation that facilitate large and small responses to a stimulus. Each brain state presented characteristic alpha oscillation features: below the critical state (*C_b_*), low amplitude was associated with an incoherent state; near the critical point (*C_p_*), high and low amplitudes were balanced with a large degree of synchronization fluctuation; above the critical point (*C_a_*), high amplitude was associated with a highly synchronized state. The alpha oscillation response was the most complex and persisted longer at critical state; in this state, responses were larger and more complex when global pulsatile stimuli were applied to a globally desynchronized, low-amplitude state. Importantly, we found there are specific phases of high-amplitude, highly synchronized alpha oscillations that barely respond to stimuli.

### Dependence on network criticality

Empirical and computational model studies have proposed that the brain in conscious wakefulness resides near a critical state, whereas brain states under significant alterations such as anesthesia and traumatic injury are distant from a critical state [28–30,36]. In this model study, we observed that the responsivity (the magnitude of significant response) at *C_b_* and *C_a_* was larger than that at *C_p_* (Fig 3D). However, the persistence of responsivity is longer at *C_p_*, which implies restricted propagation at the states far from criticality, *C_b_* and *C_a_* (Fig 3D). The brain network at *C_p_* also exhibited the most complex response when we measured perturbational complexity (Fig S1). These characteristics are consistent with TMS outcomes in empirical studies. TMS-evoked activities during unconsciousness show large but short-lasting simple response patterns [1, 3], whereas EEG responses induced by TMS during conscious wakefulness show more complex perturbational patterns compared to anesthesia and NREM sleep [1–4]. The similarity between the results of our model study and empirical results from previous studies suggests that state-dependent responses of the human brain to TMS may be due to the distinct response behaviors of the characteristic states (synchronous/incoherent or high/low amplitude) of networked oscillators below, near, and above a critical state.

### Dependence on instantaneous global synchronization

In our modeling study, we found that brain network responsiveness correlates with the level of instantaneous global synchronization at the critical state (Fig 4). Stimulation at lower levels of instantaneous global synchronization produced larger responses for both amplitude and phase synchronization, whereas higher levels of instantaneous global synchronization induced smaller responses (Fig, S12 Fig and S13 Fig). Therefore, considering the large synchronization fluctuation at a critical state, it might be important to stimulate the brain network at appropriate times in order to induce large network perturbation. Not considering the timing of stimulation may produce greater variability in outcomes. In addition, synchronization dependence provides novel insight into the role of synchronization fluctuation in brain information processing. In an epoch of lower levels of synchronization, the brain might not be able to integrate distributed information within the brain network, but may be highly susceptible to external stimuli. In contrast, in an epoch of higher levels of synchronization, the brain might easily integrate distributed information within the brain network, but may not be able to respond to external stimuli. Regarding the typical large synchronization fluctuation at a critical state, the response behaviors of a network at high and low levels of instantaneous synchronization seem to create temporal windows that can integrate internal and external information, respectively, while systematically separating internal and external information processing in the brain network.

### Dependence on the amplitude and phase of oscillation

In our modeling study, we found that responsiveness also associates with the amplitude and phase of the ongoing alpha oscillations. We decomposed the alpha oscillations into high and low instantaneous synchronization levels, amplitudes, and phases from 0° to 360°. The average responsiveness of lower instantaneous synchronization and low amplitude (LS & LA) was larger than that of higher instantaneous synchronization and high amplitude (HS & HA) (Fig 5). Importantly, considering the alpha oscillations of high synchronization and high amplitude, we found that there are specific phases (60° to 240°) that exhibit significantly smaller responsiveness in both responsiveness indexes. The results suggest periodic temporal windows (phases from 60° to 240° in each cycle of an alpha wave) in which the responsiveness of the brain network was largely inhibited.

The amplitude and phase of alpha waves (8-13Hz) are associated with sensory stimuli processing, especially in regard to visual perception [5–14]. Stimulation at lower amplitudes and at a phase near the trough of an alpha wave show increased target detectability [6,8–11,13,37]. Phosphenes artificially induced by TMS are more detectable in lower pre-stimulus alpha power and specific alpha phase ranges [7,12,38]. Recent TMS studies have also shown that motor-evoked potential (MEP) amplitude is dependent on the pre-stimulus cortical *μ*-rhythm phase [15, 16]. The established dependence of the brain’s responsiveness on the amplitude and specific phase of alpha waves is consistent regardless of stimulus type. Our human brain network model showed that stimuli applied at specific phases of alpha oscillation resulted in larger responsivity and perturbational complexity at the critical state. Interestingly, phase dependence was more pronounced when the stimulus was given at higher amplitudes of alpha oscillation (Fig 5C & 5D), which is consistent with the visual target detectability experiments [10]. This result emphasizes the importance of considering effective stimulation conditions, especially when the stimulus is applied to alpha oscillations with high amplitudes and high levels of synchronization. We also note that phase-dependent responses are diminished when stimulus strength is too small or too large (S3 Fig – S7 Fig).

We also tested if the conditions identified under global stimulation would still hold for local stimulation. We applied the same stimulus to the occipital region across three different conditions: effective (low instantaneous synchronization, low amplitude, and phases of 240°-60°), less effective (high instantaneous synchronization, high amplitude, and phases of 60°-240°), and random (conventional stimulation method without considering ongoing brain activity). Relative to the random condition, the effective and less effective conditions produced significantly larger and smaller brain responsiveness. These results may explain why the outcomes of conventional brain stimulation studies show such large inter- or intra-subject variability [39], which degrades the reliability of brain stimulation studies. In addition, finding the specific phases of alpha oscillations for large or small responsiveness reilluminates the historical concept of the “neuronic shutter” described in the 1960s [34], which implies the existence of temporal windows that periodically prohibit sensory information processing during conscious states [5,31–34,40,41]. Further study is required to identify the relevance of temporal windows of sensory information processing to the response behavior of networked alpha oscillations to external stimuli.

### Multi-scale mechanisms of large and small responsiveness

A desynchronized EEG, as characterized by low amplitude, noise, and fast-frequency activity, is associated with arousal and increased awareness [42]. This desynchronized state emerges as the product of highly recurrent and stochastic interactions within, and between, the excitatory-inhibitory neuronal populations at both the cellular and network level [42]. At the cellular level, the stochastic interplay of excitatory and inhibitory conductance sets neurons into a high-conductance state with enhanced responsiveness. At the population level, neuronal ensembles also become highly responsive to afferent inputs [42]. At the large-scale brain network level, responsiveness is determined by the characteristics of ongoing networked oscillators. However, it remains to be answered how the specific oscillation properties induce such large or small responsiveness. One of the potential mechanisms is the phase response curve (PRC) that has been studied in physics. The PRC characterizes the way that a system with a collective periodic behavior responds to external stimuli. The response of a periodically driven oscillating system is measured by the phase shift from the original phase, and the phase shift (advancing or delaying the original phase) is an inherent characteristic of any oscillatory system. This method has been applied to many rhythmic biological systems such as circadian rhythms, cardiac rhythms, and spiking neurons to study how external stimuli perturb the original rhythms [43–45]. The previous analytic studies discovered influential factors causing phase shift and phase synchronization perturbation: a low (high) phase coherence induces a large (small) phase shift [44]. Dynamically, stimulation to the phases around a stable fixed point of the PRC increases phase coherence, whereas stimulation to the phases around an unstable fixed point decreases phase coherence. These properties also hold for networks with different coupling functions, network structure, connectivity, and even for the amplitude dynamics of Stuart-Landau oscillators [46]. If the alpha oscillations are globally coupled in the human brain network [23], the response behavior may be governed by the same mechanism of PRC. Furthermore, future study needs to answer how responsiveness across multiple scales, including cellular, neural population, and large-scale brain networks, are related to each other.

## Conclusion

In summary, this modeling study demonstrated for the first time a relationship between brain network responsiveness and the ongoing alpha oscillations at different states (below, near, and above a critical state). We found properties of alpha oscillation that induce a large or small responsiveness with the same stimulus. The results explain why brain responsiveness is so variable across brain states in consciousness and unconsciousness and across time windows even during conscious wakefulness. They will also provide a theoretical foundation for developing an effective brain stimulation method that considers state-specific and transient brain activity.

## Limitations

This modeling study has several limitations. First, despite the potential contribution of other frequency bands to periodic brain responsiveness [47, 48], our model simulated only alpha oscillations. Further studies are required to confirm periodic responsiveness for other frequency bands. Our modeling results suggested that the periodic prohibition of sensory information processing in the human brain is due to the general response behavior of networked oscillations. However, to justify this argument, the association of the global synchronization of alpha oscillations (HS & HA and LS & LA) with responsiveness, which our modeling study discovered, needs to be tested empirically. Second, we only studied a pulsatile stimulus, which is similar to the stimulus type applied in previous experiments [6,8–11,13,37]. Even though we tested various stimulus strengths, we cannot assure that other types of stimulus, for instance, a continuous sinusoidal stimulus, would give the same result. Third, in this study, we mainly focused on the effect of a global stimulus and in addition tested a local stimulus targeting the occipital region. However, considering the significant influence of network topology on local brain dynamics, further study is required to identify the effects of other regional stimuli.

## Material and Methods

### Simulation of networked alpha oscillations in a human brain network structure

We constructed a large-scale functional brain network using coupled Stuart-Landau models on the human anatomical brain structure to generate spontaneous oscillations. The Stuart-Landau model with brain topology has been used to replicate the oscillatory dynamics of different types of electromagnetic brain signals such as electroencephalography (EEG), magnetoencephalography (MEG) and functional magnetic resonance imaging (fMRI) [17,18,20,24,49]. The coupled Stuart-Landau model is defined as following:

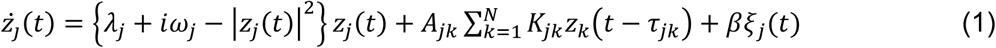

Here, the state of the *j_th_* oscillator, *j* = 1,2, …, *N*, is determined by a complex variable *z_j_*(*t*) at time *t*. N is the total number of brain regions acquired from group-averaged diffusion tensor imaging (DTI) with 82 nodes [50]. The *A_jk_* = 1 if there is a connection between the *j_th_* and *k*_*th*_ oscillators, and *A_jk_* = 0 if there is not, based on the structural brain network. Each node undergoes supercritical Hopf bifurcation and the dynamics of the oscillator settle on a limit cycle if *λ_j_* > 0, and on a stable focus if *λ_j_* < 0. Here, we fixed all *λ_j_* as 1. The *ω_j_* = 2*πf_j_* and is an initial angular natural frequency of each *j_th_* oscillator. We used a Gaussian distribution for natural frequency with a mean frequency of 10 Hz and standard deviation of 0.5 Hz to simulate the alpha bandwidth of human EEG activity [20–22,24,51]. We also used the homogeneous coupling term *K_jk_* = *K* between the *j_th_* and *k*_*th*_ oscillators from 0 to 0.4 with *δK* = 0.002, which determines the global connection strength among brain regions. The time delay between oscillators, *τ_jk_* = *D_jk_*/*s*, was introduced with the average speed of axons in brain areas, *s* = 7 *ms* [52], and the distance *Djk* between brain regions. The node *j* received input from connected node *k* after the time delay *τ_jk_*.

The time delays are various, but as long as the time delay is smaller than a quarter of the period of oscillation, here *τ_jk_* < ∼25*ms*, results are not qualitatively different [20]. A Gaussian white noise *ξ_j_*(*t*) for each node was added with the standard deviation *β* = 0.05. We numerically solved the differential equations of the Stuart-Landau model using the Stratonovich-Heun method with 1,000 discretization steps. We also tested the Runge-Kutta 4^th^ order method and the results were qualitatively same. The first 10 seconds of time series were discarded and last 25 seconds were used for the analysis of each simulation. Therefore, each brain region generates its own spontaneous oscillatory dynamics within the alpha bandwidth at each coupling strength *K* for one simulation.

### Three brain states: below, near, and above critical state

We selected three representative brain states at different coupling strengths based on the level of global synchronization and the pair correlation function (PCF) to understand the responsiveness of different systems. We first calculated an instantaneous global synchronization level *r*(*t*) at time *t* using phase difference Δθ_*jk*_(*t*) = θ_*j*_(*t*) − θ_*k*_(*t*) at each coupling strength *K*.

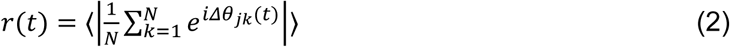

Here *r*(*t*) = 1 if all phases are equal, but *r*(*t*) is nearly zero if all phases are randomly distributed. Then we calculated the variance of *r*(*t*) for each coupling strength as a pair correlation function to define the critical state *C_p_* [53]. The pair correlation function is equivalent to susceptibility in statistical physics models. Coupled phase oscillators in complex networks showed the largest susceptibility and pair correlation function at critical state [25]. We considered generated signals at certain coupling strengths with the largest PCF to be equivalent to spontaneous alpha oscillations in conscious wakefulness. There has been emerging evidence in computational model studies that brain dynamics at critical point show the largest spatiotemporal diversity [29, 54]. Notably, a large repertoire of metastable states exists in brain dynamics during the conscious state [49]. With the critical state, we selected two other representative states, one below (*C_b_*) and one above (*C_a_*) critical state. We defined *C_b_* and *C_a_* as the states at the 10th and 90th percentiles of the averaged order parameters over all coupling strengths, respectively. These states are the representative states far from the critical states that show small PCF. Generated signals in these three states were used for the analysis. We used 20 different initial frequency distributions and selected three representative states for each frequency distribution.

### Brain network stimulation procedure

Global pulsatile stimuli were induced as a stimulation with *u*(*t*) as following:

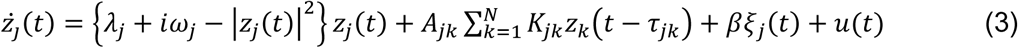

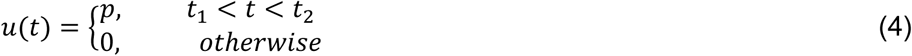

Here *p* is the strength of the stimulus during period *T* = *t*_2_ − *t*_1_. We fixed the *p* = 10 for the analysis after testing the effect of various stimulus strengths (S10 Fig.). We also fixed the duration of the stimulus, *T* = 50 *msec*, which is about half of the period of the alpha frequency range cycle. We first gave the same stimulus to the whole brain network to understand the dependence of the brain’s response derived only from the network dynamics of the system (as opposed to its network structure). The stimuli were induced at randomly selected timing *t*_1_ for one trial. In total, 30 different timings, each with 10 iterations, were selected for 20 different frequency distributions. Therefore, *C_b_*, *C_p_*, and *C_a_* each underwent 600 stimulation trials of pulsatile stimuli. Each stimulus was applied at a unique instantaneous brain state (i.e., a different one was used for each trial).

We decomposed the instantaneous brain states into the level of instantaneous global synchronization and the amplitude and phase of alpha waves to identify the relationship of each of these separate factors with responsiveness.

### Calculation of significant response after stimulation

First, we calculated the instantaneous phase synchronization of the *j*__*th*__ node, |*S*_*j*_(*t*)|, for each trial. The |*Sj*(*t*)| was obtained by taking the average of the pairwise synchronization between node *j* and node *k* at time t,

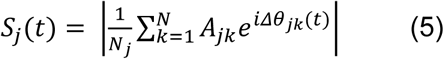

Here *N_j_* is the number of connections of the *j_th_* node. The instantaneous synchronization value for the *j_th_* node of one trial was normalized by the mean and standard deviation of the baseline synchronization values of the *j_th_* node. Baseline values were obtained by using a total of 10 seconds, consisting of 10 iterations of a 1-second pre-stimulus segment for each trial. We considered the one tail (1 − *α*) ∗ 100^*th*^ quantile with *α* = 0.05 as a significantly increased synchronization after stimulus onset. A perturbation response (PR) of the *j_th_* node at time t was defined in a binary fashion: *PR_j_*(*t*) = 1, for the significantly increased synchronization of node *j*, and *PR_j_*(*t*) = 0, otherwise.

### Responsiveness measures: responsivity and perturbational complexity

We defined two different measures to quantify the responsiveness of stimulation at different instantaneous brain states. First, a responsivity *R_j_* was defined by taking the average *PR_j_* during a specific epoch after stimulation. It quantifies the total amount of significant responses after stimulation. Here we used t = 500 msec, which covers the maximum response changes after stimulation.

Second, we also calculated the Lempel-Ziv complexity (LZc) of the PR to understand how complex the response was. The LZc of the *PR_j_* was defined as following:

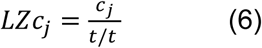

Here (*c_j_*) is the non-normalized Lempel-Ziv complexity of *PR_j_* calculated by a LZ76-algorithm [35], i.e., the number of unique patterns in the *PR_j_* during time *t*. We defined a perturbational complexity, *LZc_j_* as the normalized *c_j_*.

We first investigated the relationships between the level of instantaneous global synchronization at the stimulus onset *r_s_*, *s* = 1,2, …, 600, and the average *R* and *LZc* over all nodes at *C_b_*, *C_p_*, and *C_a_* (Fig 4). Here we calculated spatial *LZc*, which is 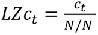, *N* = 82 to see the global effect *N*/*N* of the stimulation. Spearman correlations with p-value were calculated between *r_s_* and the average *R* and *LZc* at *C_b_*, *C_p_*, and *C_a_*. A Kruskal-Wallis test with a multiple comparison test using the Tukey-Kramer method was used to see the statistical differences across brain states (***p<0.001).

We also examined the instantaneous alpha amplitude/phase dependences of responsiveness (*R* and *LZc*) to the stimuli (Fig 5). Here we found the certain oscillatory conditions of alpha oscillations that can induce large or small responsiveness at *C_p_*. We considered the oscillation properties with certain levels of instantaneous global synchronization, amplitudes, and phases that induce large (small) responsiveness as effective (less effective) stimulation conditions and applied these conditions to stimulate the local areas described below.

The same procedures were performed for the amplitude response; these results are shown in the supplementary materials (S2 Fig – S13 Fig).

### Local stimulation effect

At *C_p_*, we induced stimuli to 13 regions in the occipital area within a 50 mm radius of the left cuneus to confirm that the specific stimulation conditions we found from global stimulation could be applied to local stimulation (Fig 6). On the left and right hemispheres of the brain, respectively, the stimulated regions included: (left) cuneus, inferior parietal, isthmus cingulate, lateral occipital, lingual, pericalcarine, precuneus, superior parietal; (right) cuneus, isthmus cingulate, lingual, pericalcarine, precuneus. We selected the occipital region because abundant previous sensory stimulation studies mostly targeted the occipital region for visual tasks within the alpha frequency band. We compared the responsiveness measures with effective, less effective and random stimulation conditions. For the effective (less effective) stimulation, we applied the stimulus to the alpha phases from 240° to 60° (60° to 240°) with low (high) levels of instantaneous global synchronization and low (high) amplitude. The stimuli were applied to the random instantaneous brain states of the occipital region for the random stimulation condition. The number of nodes for random stimulation was determined by a mean value of the number of nodes for effective and non-effective conditions. We performed 600 different trials at *C_p_*, in which each trial consisted of stimuli with *p* = 30 for 50 msec, and classified the target nodes with specific conditions. We used a Kruskal-Wallis test to compare the significant differences in responsiveness for three conditions. A multiple comparison test was also performed using the Tukey-Kramer method (**p<0.005 and ***p<0.001).

## Acknowledgments

This work is supported by grant No. R01 GM098578 (PIs: GM and UL) from the National Institutes of Health.

