## Supplementary data for "Alpha oscillation, criticality, and responsiveness in complex brain networks"

**S1 Fig.**

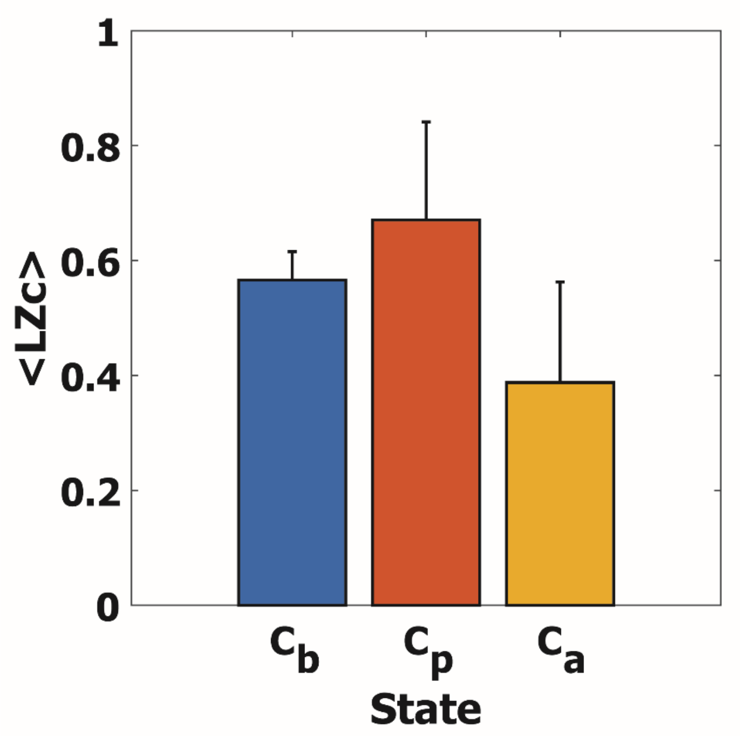


**S1 Fig. Average perturbational complexity LZc over all trials for $C_b$, $C_p$, and $C_a$.** The error bar indicates a standard deviation. Average LZc is the largest at $C_p$ (mean±SD = 0.57±0.05, 0.67±0.17, and 0.39±0.18 for $C_b$, $C_p$, and $C_a$), indicating the most complex and flexible responses exist at $C_p$.

In the supplementary material, we show responsiveness indexes of both amplitude and phase synchronization because it has been reported that the amplitude and phase dynamics have different criticality features [1]. Therefore, we additionally investigated a perturbation response of the amplitude, which is also thought like the power change after the stimulation in previous empirical experiments. As explained in the main text, responsivity and perturbational complexity of the amplitude response ($\mathrm{R}^{\mathrm{amp}}$ and $\mathrm{LZc}^{\mathrm{amp}}$) were calculated in the same way of the synchronization response ($\mathrm{R}^{\mathrm{sync}}$ and $\mathrm{LZc}^{\mathrm{sync}}$), but with the 1500 msec epochs after the stimulation.


1. Daffertshofer A, Ton R, Kringelbach ML, Woolrich M, Deco G. Distinct criticality of phase and amplitude dynamics in the resting brain. Neuroimage. Elsevier Ltd; 2018; 1–6. doi:10.1016/j.neuroimage.2018.03.002

### S2 Fig. Stimuli strength p = 10

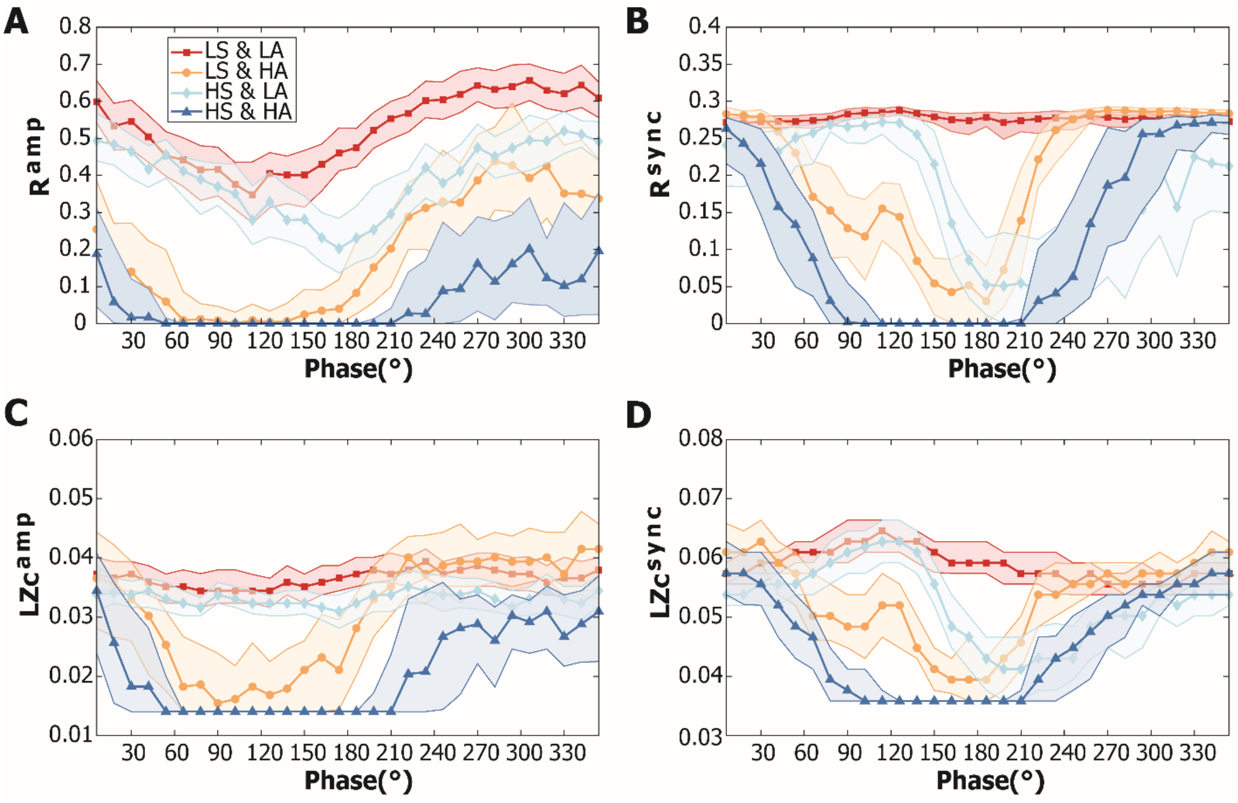


**S2 Fig. Responsiveness of nodes with different stimulation conditions at $C_p$.** For each panel, 4 different conditions were classified; LS & LA (red), LS & HA (orange), HS & LA (light blue), and HS & HA (blue). LS (LA) indicates a globally desynchronized (low amplitude) state at stimulus onset. HS (HA) indicates a globally synchronized (high amplitude) state at stimulus onset. Lower and higher levels of instantaneous global synchronization (amplitude) were distinguished by the average of instantaneous global synchronization levels across all trials (average of amplitudes across all nodes). See the main text for more detailed information. Colored thick lines indicate the median (A) $R^{amp}$, (B) $R^{sync}$, (C) $LZc^{amp}$, (D) $LZc^{sync}$ of nodes. Shaded areas cover the 40% to 60% quantiles of each responsiveness. LA conditions produce large amplitude responsiveness. HA conditions appear to produce smaller responsiveness in phases 60° to 240°.

### S3 Fig. Stimuli strength p = 2

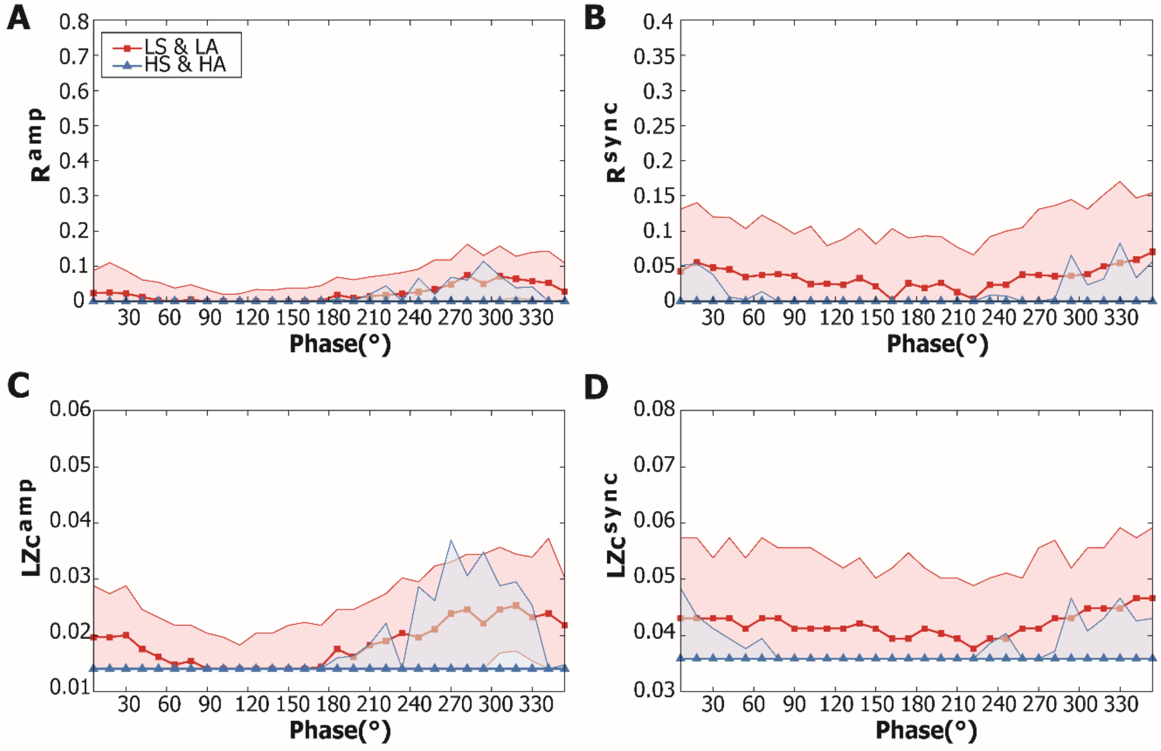


**S3 Fig. Responsiveness of nodes with different stimulation conditions at $C_p$ for the stimuli strength $p = 2$.** For each panel, a red (blue) thick line indicates the median (A) $R^{amp}$, (B) $R^{sync}$, (C) $LZc^{amp}$, (D) $LZc^{sync}$ of nodes with LS&LA and HS&HA state at stimulus onset. Lower and higher levels of instantaneous global synchronization (amplitude) were distinguished by the average of instantaneous global synchronization levels across all trials (average of amplitudes across all nodes). Phases of nodes at stimulus onset were sorted in 30 phase bins. Shaded areas cover the 25% to 75% quantiles of each responsiveness. Responsiveness is small for all measures.

**S4 Fig. Stimuli strength p = 5**

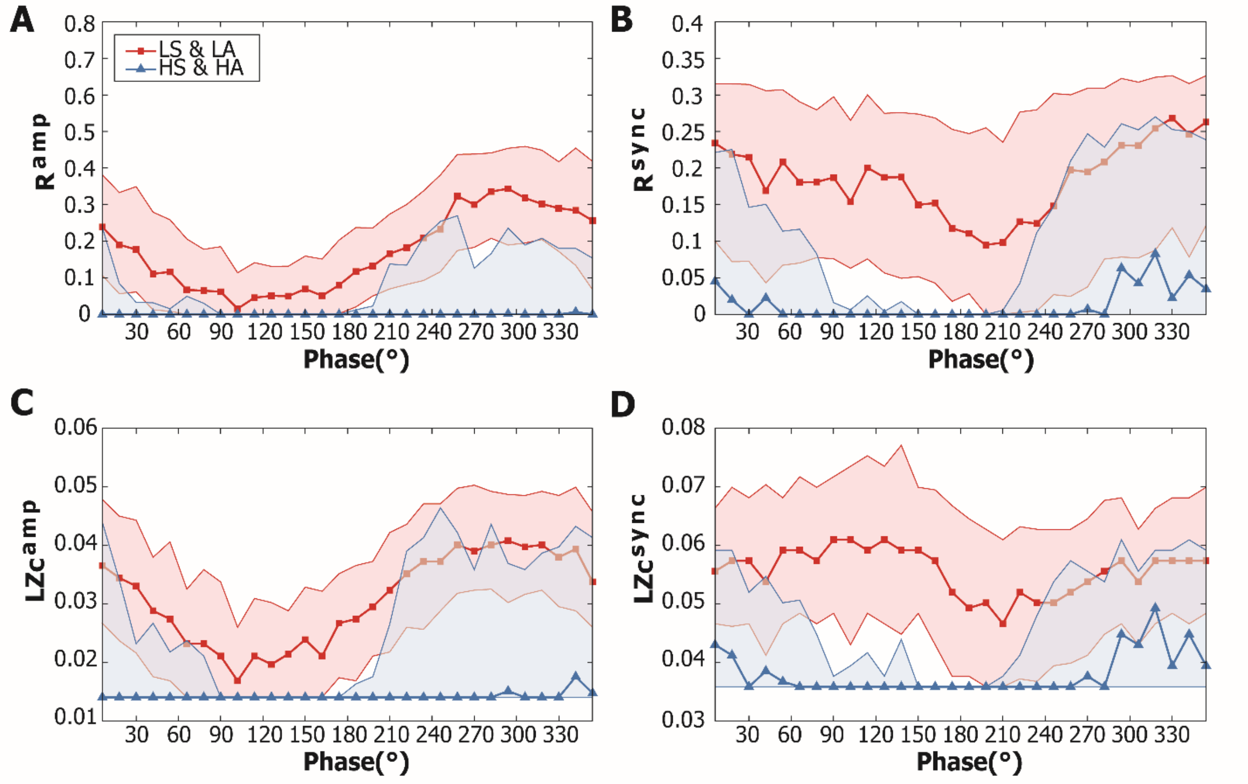


**S4 Fig. Responsiveness of nodes with different stimulation conditions at $C_p$ for the stimuli strength $p = 5$.** For each panel, a red (blue) thick line indicates the median (A) $R^{amp}$, (B) $R^{sync}$, (C) $LZc^{amp}$, (D) $LZc^{sync}$ of nodes with lower (higher) level of instantaneous global synchronization and lower (higher) amplitude at stimulus onset. Lower and higher levels of instantaneous global synchronization (amplitude) were distinguished by the average of instantaneous global synchronization levels across all trials (average of amplitudes across all nodes). Phases of nodes at stimulus onset were sorted in 30 phase bins. Shaded areas cover the 25% to 75% quantiles of each responsiveness. Phase dependences are shown in all four responsiveness measures.

### S5 Fig. Stimuli strength p = 20

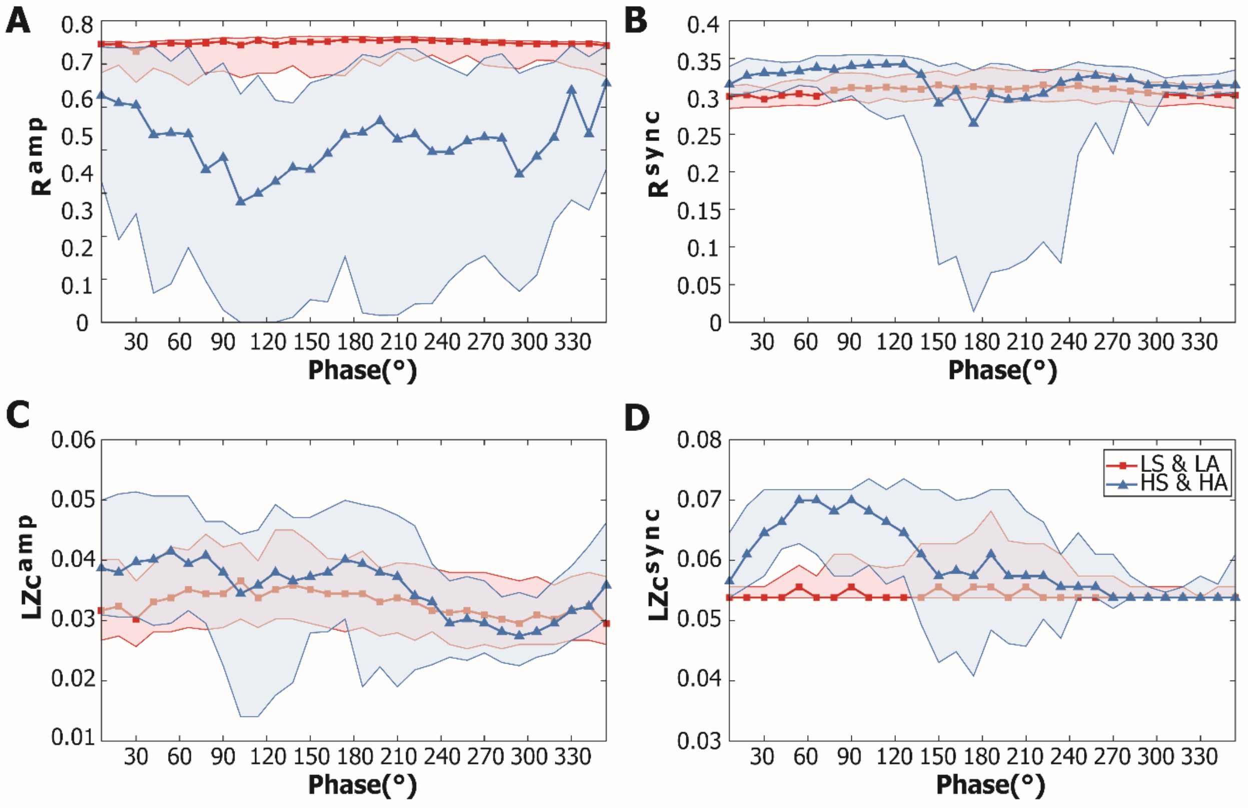


**S5 Fig. Responsiveness of nodes with different stimulation conditions at $C_p$ for the stimuli strength $p = 20$.** For each panel, a red (blue) thick line indicates the median (A) $R^{amp}$, (B) $R^{sync}$, (C) $LZc^{amp}$, (D) $LZc^{sync}$ of nodes with lower (higher) level of instantaneous global synchronization and lower (higher) amplitude at stimulus onset. Lower and higher levels of instantaneous global synchronization (amplitude) were distinguished by the average of instantaneous global synchronization levels across all trials (average of amplitudes across all nodes). Phases of nodes at stimulus onset were sorted in 30 phase bins. Shaded areas cover the 25% to 75% quantiles of each responsiveness. Amplitude responsivity ($R^{amp}$) of LS&LA seems large, but the dependence of oscillation properties at the stimulus onset appear to diminish in complexity measures.

**S6 Fig. Stimuli strength p = 30**

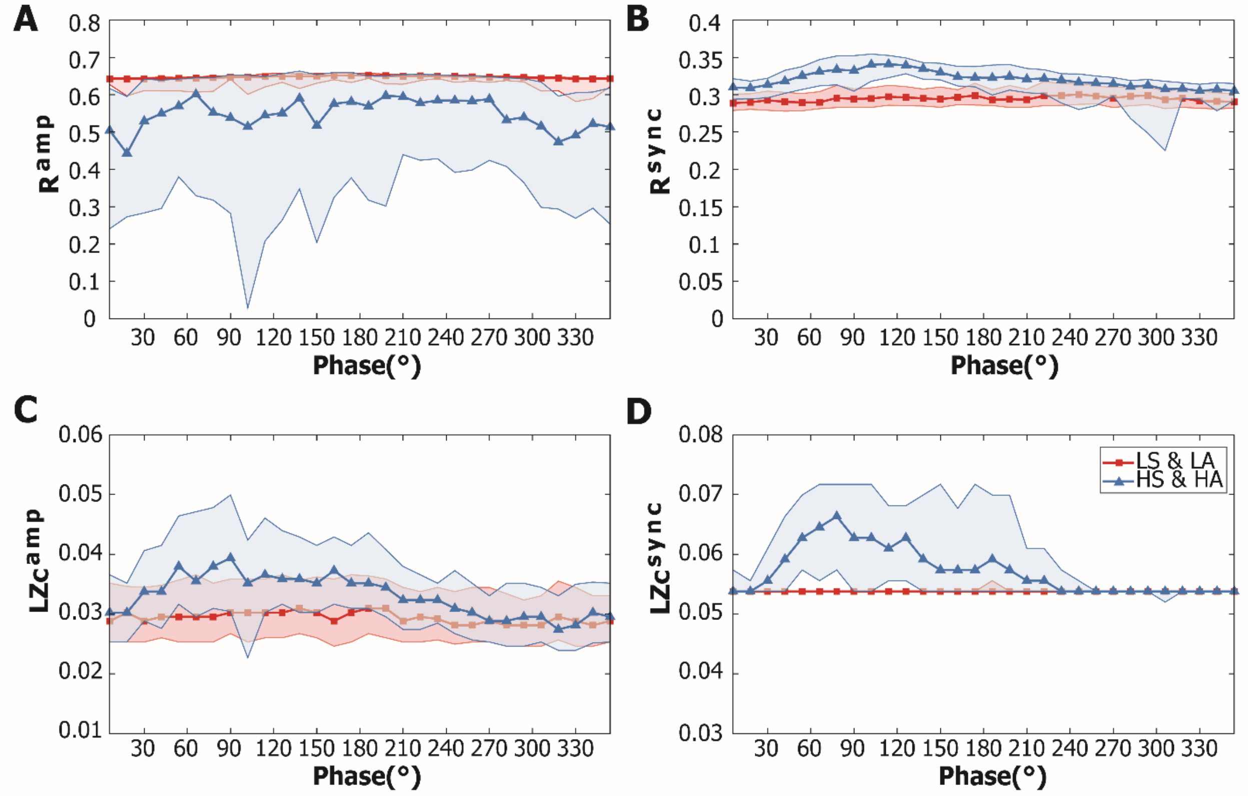


**S6 Fig. Responsiveness of nodes with different stimulation conditions at $C_p$ for the stimuli strength $p = 30$.** For each panel, a red (blue) thick line indicates the median (A) $R^{amp}$, (B) $R^{sync}$, (C) $LZc^{amp}$, (D) $LZc^{sync}$ of nodes with lower (higher) level of instantaneous global synchronization and lower (higher) amplitude at stimulus onset. Lower and higher levels of instantaneous global synchronization (amplitude) were distinguished by the average of instantaneous global synchronization levels across all trials (average of amplitudes across all nodes). Phases of nodes at stimulus onset were sorted in 30 phase bins. Shaded areas cover the 25% to 75% quantiles of each responsiveness. Amplitude responsivity ($R^{amp}$) of LS&LA seems large, but the dependence of oscillation properties at stimulus onset appear to diminish in the other three measures.

### S7 Fig. Stimuli strength p = 40

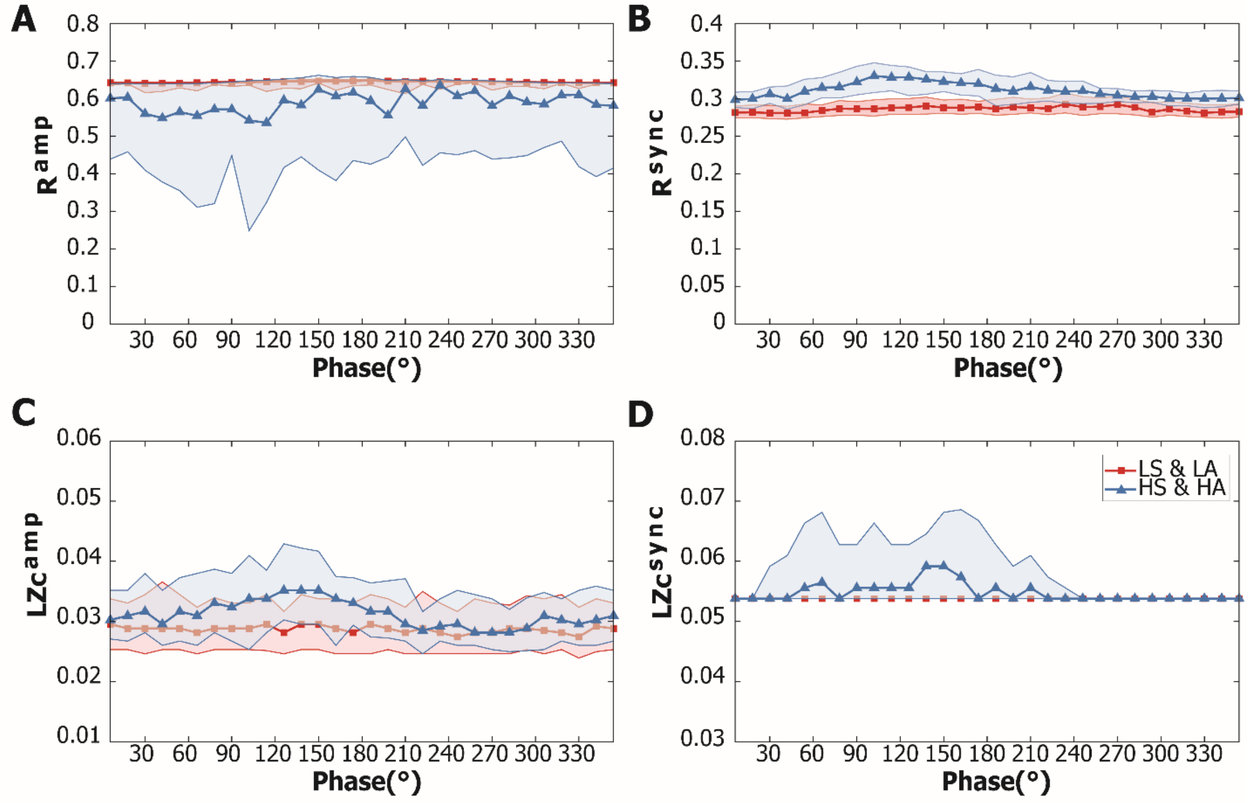


**S7 Fig. Responsiveness of nodes with different stimulation conditions at $C_p$ for the stimuli strength $p = 40$.** For each panel, a red (blue) thick line indicates the median (A) $R^{amp}$, (B) $R^{sync}$, (C) $LZc^{amp}$, (D) $LZc^{sync}$ of nodes with lower (higher) level of instantaneous global synchronization and lower (higher) amplitude at stimulus onset. Lower and higher levels of instantaneous global synchronization (amplitude) were distinguished by the average of instantaneous global synchronization levels across all trials (average of amplitudes across all nodes). Phases of nodes at stimulus onset were sorted in 30 phase bins. Shaded areas cover the 25% to 75% quantiles of each responsiveness. Amplitude responsivity ($R^{amp}$) of LS&LA seems large, but the dependence of oscillation properties at the stimulus onset appear to diminish in all measures.

### S8 Fig.

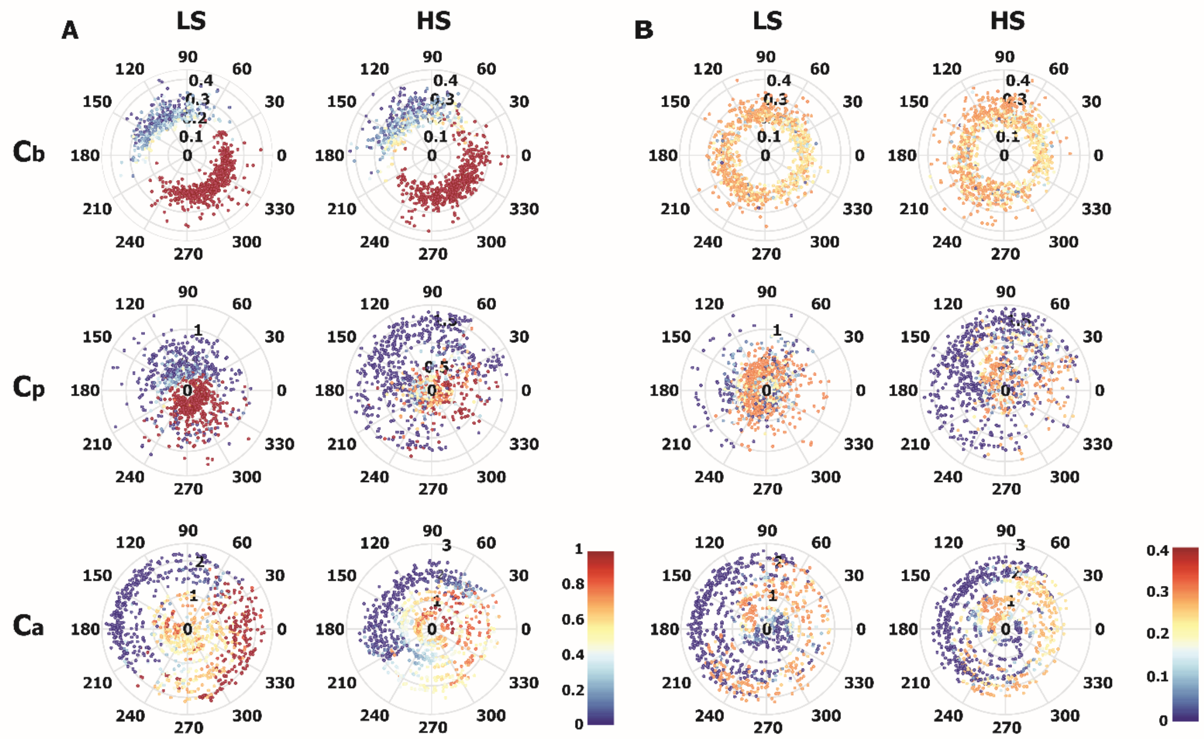


**S8 Fig. Instantaneous global synchronization level, node amplitude and phase dependences of responsivity $\boldsymbol{R}$ at $\boldsymbol{C_b}$, $\boldsymbol{C_p}$, and $\boldsymbol{C_a}$.** (A) Sixty trials that stimulation was given at the lowest (left) / highest (right) levels of instantaneous global synchronization (LS and HS, respectively) were first selected for $C_b$, $C_p$ and $C_a$ (first, second, and third row). Dots indicate 10% of nodes with the largest (smallest) $R^{amp}$ within those trials. Color of dots indicates the value of $R^{amp}$ of the node. A distance from the origin of the dot is the node amplitude, and a phase angle of the dot is the node phase at stimulus onset. The phase dependence is prominent for $R^{amp}$ at $C_b$. Stimulating at phase ranges from 60° to 240° of the oscillation produces smaller $R^{amp}$ (bluish) for all three states. (B) Color of dots indicates the value of $R^{sync}$ of the node. Stimulating at lower amplitude (shorter distance from the origin) tend to show larger $R^{sync}$ (reddish) in HS at $C_p$. Stimulating at lower amplitude tend to show larger $R^{sync}$ at $C_a$.

**S9 Fig.**

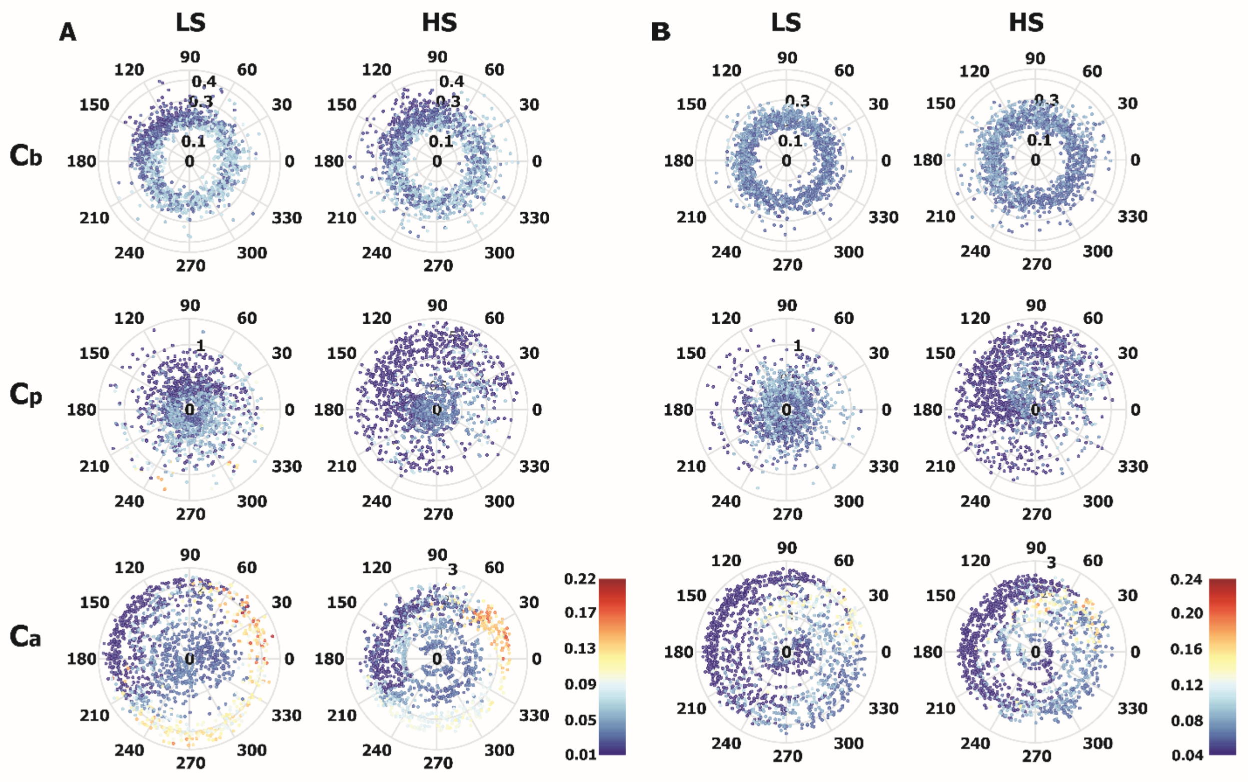


**S9 Fig. Instantaneous global synchronization level, node amplitude and phase dependences of perturbational complexity $LZc$ at $C_b$, $C_p$, and $C_a$.** (A) Sixty trials that stimulation was given at the lowest (left) / highest (right) levels of instantaneous global synchronization (LS and HS, respectively) were first selected for $C_b$, $C_p$ and $C_a$ (first, second, and third row). Dots indicate 10% of nodes with the largest (smallest) $LZc^{amp}$ within those trials. Color of dots indicates the value of $LZc^{amp}$ of the node. A distance from the origin of the dot is the node amplitude, and a phase angle of the dot is the node phase at stimulus onset. The phase dependence is shown in $LZc^{amp}$ at $C_b$. Stimulating at lower amplitude (shorter distance from the origin) tend to show larger $LZc^{amp}$ at $C_p$ in HS. Stimulating at phase ranges from 240° to 60° at $C_a$ shows larger $LZc^{amp}$ (reddish). (B) Color of dots indicates the value of $LZc^{sync}$ of the node. Stimulating at lower amplitude tend to have larger $LZc^{sync}$ in HS at $C_p$. Stimulating at phases from 240° to 60° at $C_a$ shows larger $LZc^{sync}$.

**S10 Fig.**

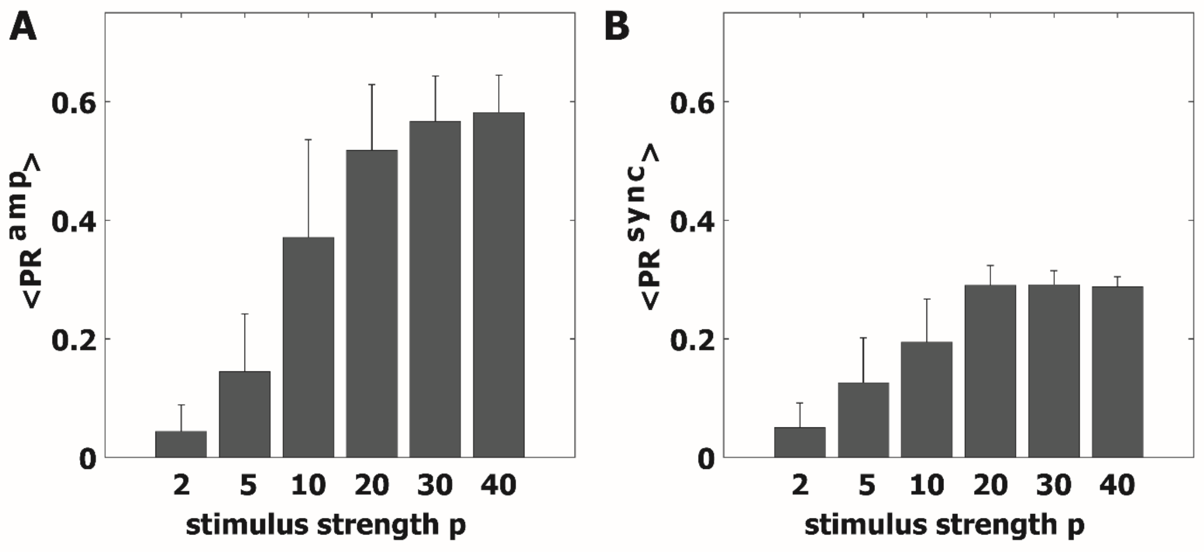


**S10 Fig. Average (A) $PR^{amp}$ and (B) $PR^{sync}$ over all trials with respect to the pulsatile stimuli strength $p$.** The error bar indicates a standard deviation. $< PR^{amp} >$ is increased when $p$ is increased and $< PR^{sync} >$ is increased when $p$ is increased until the strength=30.

**S11 Fig.**

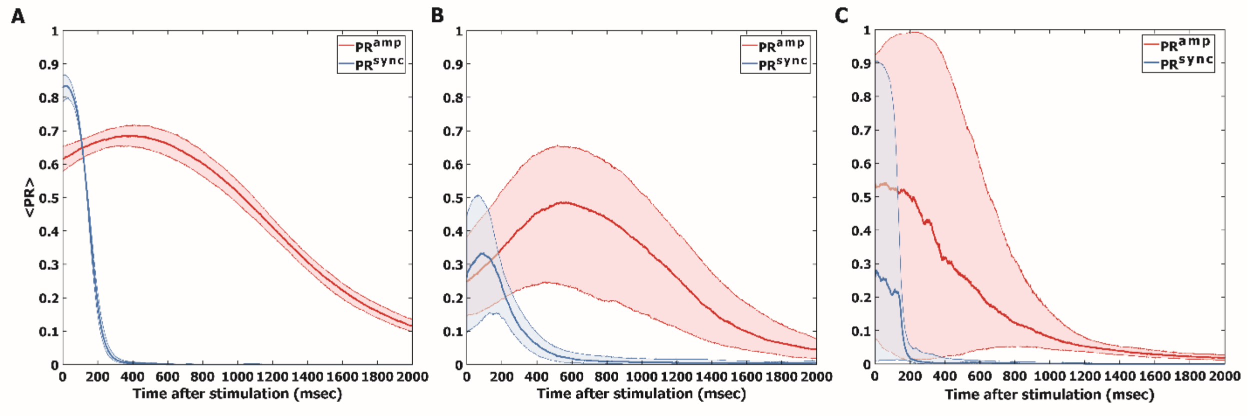


**S11 Fig. Average $PR^{amp}(t)$ and $PR^{sync}(t)$ over all trials for three brain states.** Average $PR^{amp}(t)$ and $PR^{sync}(t)$ at (A) $C_b$, (B) $C_p$, and (C) $C_a$. Thick lines indicate the median of the average PR across all trials and shaded areas cover the 25% to 75% quantiles. $PR^{amp}$ and $PR^{sync}$ are differentiable, that is, the $PR^{amp}$ maintains longer than the $PR^{sync}$, implying that there might be different criticality features in amplitude and phase dynamics [1].

**S12 Fig.**

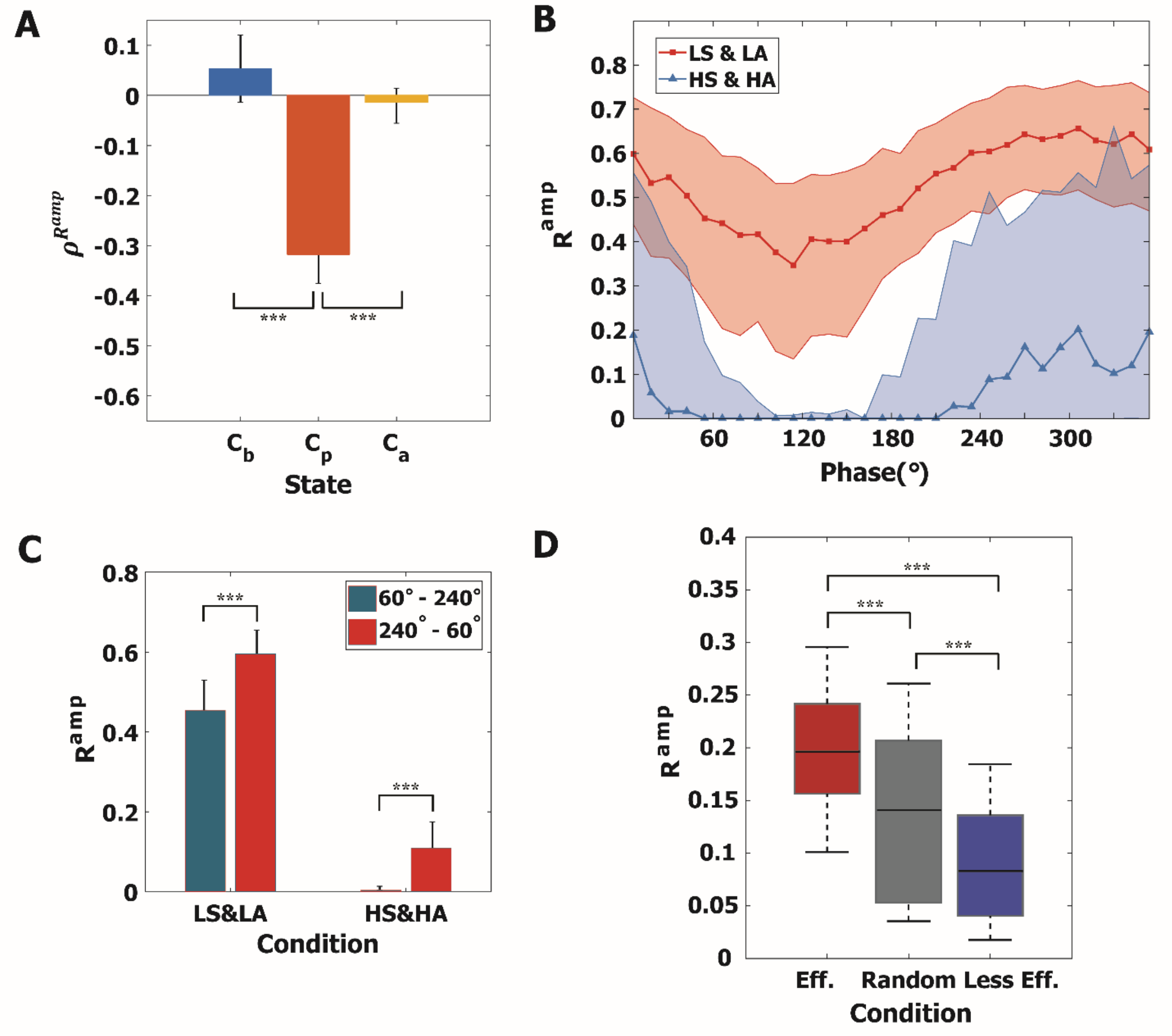


**S12 Fig. Responsivity results from the amplitude response.** The amplitude responsivity $R^{amp}$ shows similar results with the synchronization responsivity $R^{sync}$. (A) Negative Spearman correlation between $r_s$ and $R^{amp}$ (-0.33). (B) $R^{amp}$ of nodes with different stimulation conditions at $C_p$. Stimulating at LS&LA (red) shows larger responsiveness to the stimulus than stimulating at HS&HA (p<0.001, multiple comparison test using Tukey-Kramer method). (C) $R^{amp}$ of 4 representative stimulation conditions. For HS & HA, stimulation at specific phases of alpha waves from 60° to 240° shows significantly smaller responsivity (***p<0.001, Wilcoxon rank sum test). (D) $R^{amp}$ of the local stimulation with effective, random, and less effective stimulation conditions. The effective and less effective stimulation conditions we found in the global stimulation hold for local stimulation.

**S13 Fig.**

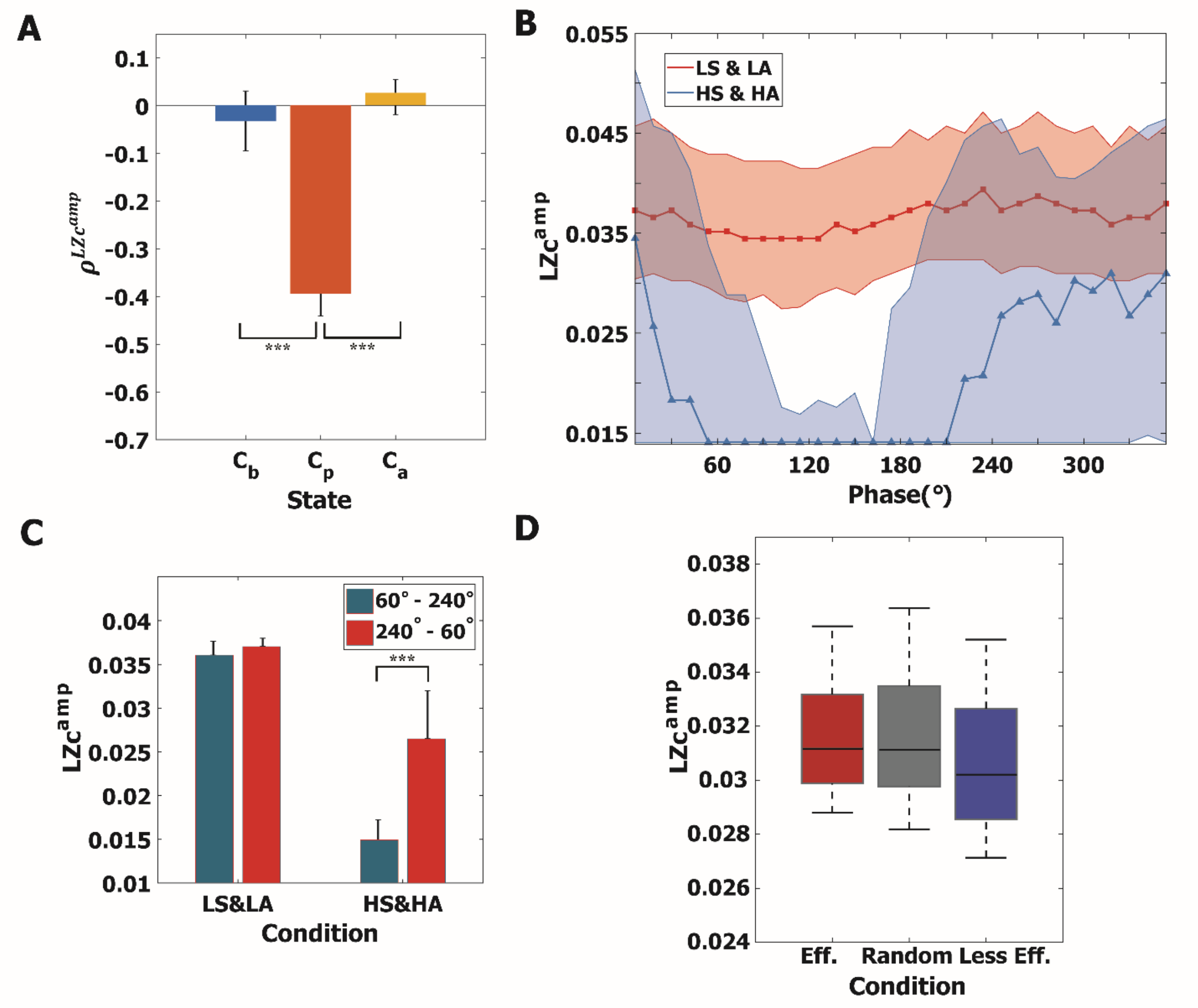


**S13 Fig. Perturbational complexity results from the amplitude response.** The amplitude complexity $LZc^{amp}$ shows similar results with the synchronization complexity. (A) Negative Spearman correlation between $r_s$ and $LZc^{amp}$ (-0.39). (B) $LZc^{amp}$ of nodes with different stimulation conditions at $C_p$. Stimulating at LS&LA (red) shows larger responsiveness to the stimulus than stimulating at HS&HA (p<0.001, multiple comparison test using Tukey-Kramer method). (C) $LZc^{amp}$ of 4 representative stimulation conditions. For HS & HA, stimulation at specific phases of alpha waves from 60 ° to 240 ° shows significantly smaller complexity (***p<0.001, Wilcoxon rank sum test). (D) $LZc^{amp}$ of the local stimulation with effective, random, and less effective stimulation conditions. $LZc^{amp}$ shows no significant difference across conditions, which is different from the results of synchronization complexity. It might be related to effects from the network structure during the long duration after the stimulation.
